# Boron induces osteogenesis by stimulating NaBC1 in cooperation with BMPR1A

**DOI:** 10.1101/2020.03.17.995001

**Authors:** Patricia Rico, Aleixandre Rodrigo-Navarro, Laura Sánchez Pérez, Manuel Salmeron-Sanchez

## Abstract

The intrinsic properties of Mesenchymal Stem Cells (MSCs) make them ideal candidates for tissue engineering applications as they are regulated by the different signals present in the stem cell niche. Considerable efforts have been made to control stem cell behavior by designing material system approaches to engineer synthetic extracellular matrices and/or include soluble factors in the media. This work proposes a novel and simple approach based on ion-channel stimulation to determine stem cell fate that avoids the use of growth factors (GFs). We used boron ion - essential item in cell metabolism - transported inside cells by the NaBC1-channel. Addition of boron alone enhanced MSC adhesion and contractility, promoted osteogenesis and inhibited adipogenesis. The stimulated NaBC1 promoted osteogenesis via activation of the BMP canonical pathway (comprising Smad1 and YAP nucleus translocation and osteopontin expression) through a mechanism that involves simultaneous NaBC1/BMPR1A and NaBC1/α_5_β_1_/α_v_β_3_ co-localization,. We describe a novel function for NaBC1 as a mechanosensitive ion-channel capable of interacting and stimulating GF receptors and fibronectin-binding integrins. Our results open up new biomaterial engineering approaches for biomedical applications by a cost-effective strategy that avoids the use of soluble GFs.

## 1. Introduction

Mesenchymal stem cells (MSCs) are multipotent and capable of differentiating mesodermal lineages (reticular, adipogenic, osteogenic and chondrogenic) under certain conditions that often include growth factors (GFs)^[1]^. Their final fate *in vivo* depends on a combination of physical, chemical and biological cues, all of which are present in the natural stem cell niche. The extracellular matrix (ECM) of the stem cell niche plays an important role by supporting cell growth, dynamically regulating GF release and activating multiple intracellular pathways^[2,3]^. Material systems offer an alternative for accurately engineering synthetic ECMs. The current material-based strategies for directing stem cell behavior are based on varying the material properties in terms of chemistry^[4]^, stiffness^[5,6]^ or topography^[7,8]^. Other approaches use GFs dissolved in the media, delivered from material systems or as components of bioactive materials for efficient solid-phase presentation^[9]^. However, although they are widely used clinically in high concentrations, they can also produce undesired side effects such as neurological problems or cancer^[10]^, so that alternative methods are needed to avoid supraphysiological doses of GFs.

Among the many factors governing MSCs commitment, mechanical cues have emerged as key determinants in cell fate and function in a context-dependent manner^[11]^. Force exerted by the cells on the ECM is mediated by major adhesion receptors, integrins, mechanosensors that pull on ECM proteins and transmit mechanical forces. This interaction causes cytoskeleton contraction and integrin clustering, giving rise to focal adhesions^[12]^ and an intracellular cascade of downstream signal transduction events that determine the stem cell fate. It is widely accepted that the osteogenic/adipogenic balance is determined by the initial MSC adhesion: i.e. poorly adhered MSCs (small focal adhesions) differentiate into rounded adipocytes^[7,13]^ while MSCs that develop mature adhesions (long in size, allowing cell spreading and intracellular tension) differentiate into osteoblasts^[6,13]^. Besides integrins, cells can also perceive force through ion-channels. This activation occurs after ligand binding, membrane stretch, interaction with other specific ligands or changes in membrane potential^[14]^, so that ion-channels can act as mechanosensors that communicate extracellular signals to the cytoplasmic environment^[15]^ and integrins^[16]^. The contribution of ion-channels to the regulation of cell behavior in MSCs is becoming increasingly recognized, although the precise mechanisms are still debated and the information available mainly refers to the Ca^2+^, K^+^ and Cl^-^ channels^[17]^.

The NaBC1-channel controls boron (B) homeostasis and, in the presence of borate, functions as an obligated Na^+^-coupled borate co-transporter^[18]^. Mutations in the NaBC1 gene cause endothelial corneal dystrophies^[19,20]^ and are overexpressed in most breast cancer cell lines and downregulated in some colorectal tumors^[18]^. We previously reported that B promotes myogenic differentiation^[21]^, proposed a 3D molecular model for NaBC1 and showed that the simultaneous stimulation of NaBC1 and the vascular endothelial growth factor receptor (VEGFR) promote angiogenesis *in vitro* and *in vivo* with ultralow-doses of GFs^[22]^. Although little is known about boron homeostasis and function in mammalian cells, there are several reports that describe the role of this metalloid-enhancing MSC osteogenic differentiation^[23-25]^, and only recently others have described the inhibition of adipogenic differentiation^[26,27]^.

Here we report that addition of boron to the culture media induces osteogenesis and inhibits adipogenesis in the absence of other soluble chemicals or GFs. We propose a novel mechanism involving crosstalk and co-localization of active NaBC1/BMPR1A and NaBC1/α_5_β_1_/α_v_β_3_ integrins that activate intracellular pathways, with NaBC1 in the novel function of a mechanosensitive ion-channel.

## 2. Results

With the aim of evaluating MSC osteogenic commitment, we used a combined system for simultaneous stimulation of NaBC1 and α_5_β_1_ and α_v_β_3_ integrins^[28]^. Boron (B) in the form of Sodium Tetraborate Decahydrate (borax) was used dissolved in the culture media for NaBC1 stimulation, and fibronectin (FN) for FN-binding integrin activation. We used Glass (control substrate) and polylactic acid (PLLA) as a possible FDA approved^[29]^ biodegradable substrate for B delivery in future *in vivo* applications.

We used C3H10T1/2 cells, a murine multipotent MSC line from mouse embryo as the model cellular system, widely used in the study of cell differentiation mechanisms and capable of differentiation into mesodermal lineages (reticular, adipogenic, osteogenic and chondrogenic) ^[30,31]^.

We first evaluated boron cytoxicity to assess the maximum working concentrations for this particular cell line. **Figure S1** shows the viability results obtained, indicating that boron concentrations higher than 26.2 mM (10 mg mL^-1^) are toxic for C3H10T1/2 cells. In all the experiments, we supplemented the culture media with 0.59 mM (labeled B2%) and 1.47 mM (labeled B5%) from a B solution, concentrations within the 0.2 - 0.6 mg mL^-1^ range, to avoid any boron toxic effects on cells.

### 2.1. NaBC1 stimulation induces MSCs adhesion

All the substrates employed were coated with a 20 µg mL^-1^ concentration of human plasma FN solution. We previously reported that FN adsorbs similarly on Glass and PLLA substrates in the presence/absence of B, disregarding any influence of the ion on FN surface density^[21]^.

We evaluated the combined effect of stimulating NaBC1 together with FN-binding integrins after 3 h of culture using serum-depleted culture media to ensure that initial cell-material interaction was exclusively FN-mediated, and so targeting α_5_β_1_ and α_v_β_3_ integrins^[28]^. Cells were seeded at low density (5,000 cells cm^-2^) to favor cell-material interaction and minimize cell-cell contacts in order to quantify focal adhesions (FA). We have performed the experiments using intact (non-transfected cells), transfected MSCs with esiRNA^NC^ (fluorescent-labeled universal negative control with no sequence homology to any known gene sequence) and esiRNA^NaBC1^ (with specific NaBC1 sequence homology). We first assess the transfection efficiency of the silencing experiment, detecting a clear red fluorescence in MSC cells after 24 h and 3 days post-transfection (**Figure S2**-a). Real time quantitative PCR (qPCR) amplification resulted in a decrease of NaBC1 mRNA levels after transfecting cells with esiRNA^NaBC1^, confirming a successful silencing of the transporter (Figure S2-b).

Considering the stiffness range of the Glass (50-90 GPa)^[32]^ and PLLA (3.5 GPa)^[33]^ used as substrates, and the range of force that cells can exert (up to 5 nN µm^-2^)^[34]^, we hypothesized that the cells would detect all the substrates as rigid and so no result can be attributed to differences in the mechanical sensing of the environment. **Figure 1-a** shows MSC morphology before and after NaBC1 silencing. Cells displayed marked actin stress fibers ending at well-developed focal adhesion sites of attachment that were more evident in the PLLA-B2% and PLLA-B5% substrates in non-transfected and esiRNA^NC^ cells. Quantification of cell parameters showed that the presence of boron induced significant differences in the cell spreading area, even though the same number of cells was used in all the substrates (Figure S2-c).

**Figure 1.**
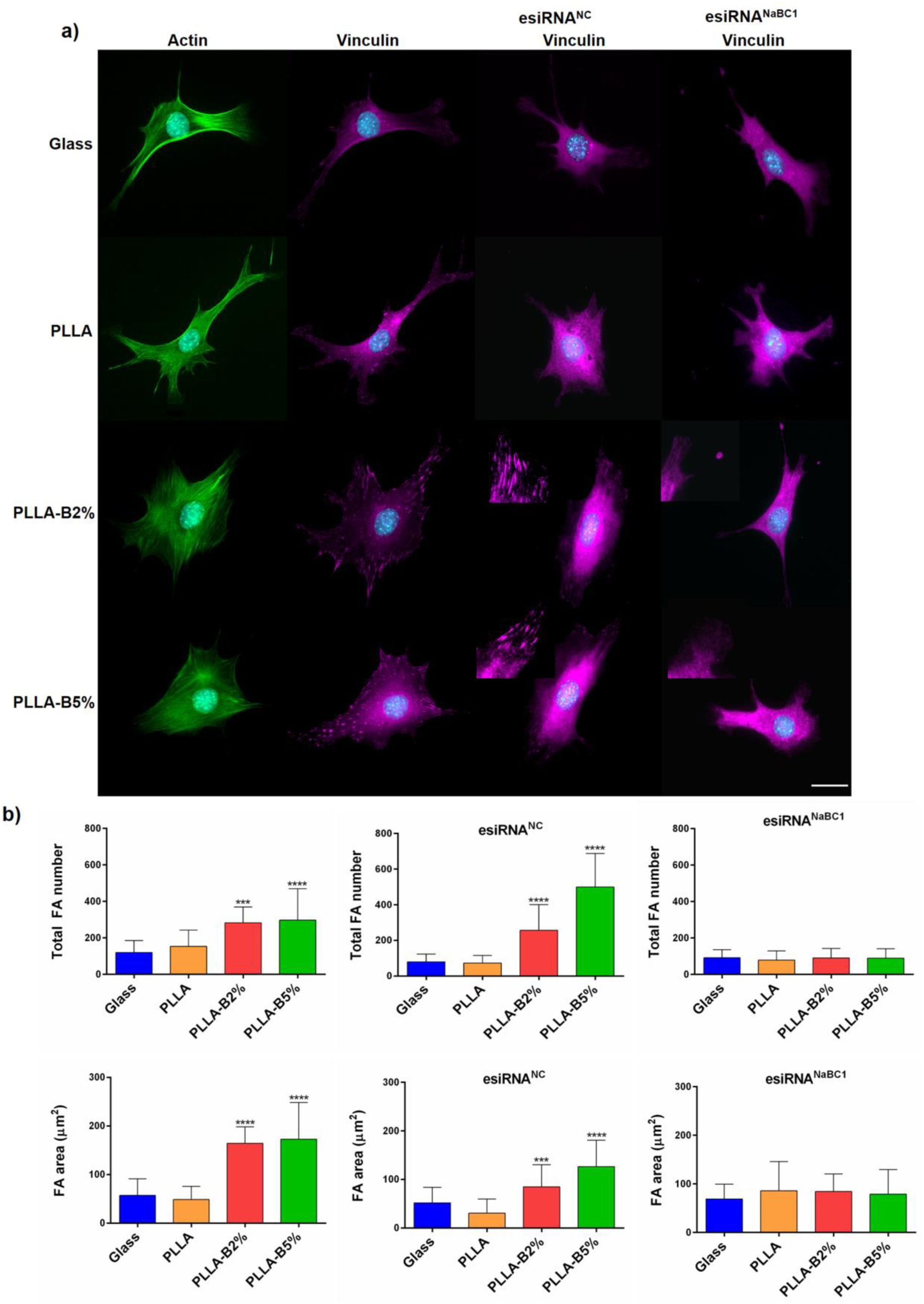
Boron effects on Focal Adhesion (FA) formation and MSC adhesion. a) Immunofluorescence images of actin cytoskeleton (green), nuclei (cyan) and vinculin (magenta) as a marker of focal adhesions (FA). MSCs were cultured for 3 h, onto functionalized substrates (FN-coated), with serum-depleted and boron (PLLA-B2%, PLLA-B5%) presence in the culture medium. The same experiment was performed after NaBC1 silencing using esiRNAs (esiRNA^NaBC1^). Universal negative control esiRNA (esiRNA^NC^) was used as a transfection efficiency control. The PLLA-B2% and PLLA-B5% substrates presented clear focal adhesions as a result of vinculin staining that strongly diminished after NaBC1 silencing (see inset magnifications). Scale bar 25 µm. b) Image analysis quantification of different parameters related to FA. Total FA number and FA area before and after NaBC1 silencing. Boron induces more and bigger FAs, effect that is reverted after NaBC1 silencing. Statistics are shown as mean ± standard deviation. n = 20 images/condition from three different biological replicas. Data was analyzed by an ordinary one-way ANOVA test and corrected for multiple comparisons using Dunnett analysis (P = 0.05). ***p < 0.001, ****p < 0.0001.

FA quantification gave a higher total FA number and larger FA area in the presence of B (Figure 1-b). After NaBC1 silencing, we observed a strong decrease in the total FA number and area, even in the presence of B (Figure 1-a and 1-b), indicative that the enhancement of MSC adhesion depends on a functional NaBC1. The FA distribution frequency was similar on all the surfaces, with a greater number of nascent plaques (between 0-6 µm), which decreased monotonically until they became mature plaques (> 6 µm). A detailed analysis of FA distribution revealed that boron induced FA formation at different FA development stages, resulting in higher levels of nascent (0-6 µm) and mature FA (6-12 µm) (Figure S2-d). Since integrins initiate the adhesion process by generating nascent integrin-matrix linkages that afterwards develop into mature integrin-ECM linkages, recruiting additional components under force^[35]^, our results indicate that active NaBC1 accelerates integrin clustering to form FA in the early stages that will become mature FA in PLLA-B2% and PLLA-B5% substrates, and demonstrates that NaBC1 activation regulates cell adhesion as previously reported for other ion-channels^[36]^.

We next examined whether cell adhesion triggered by boron activation of NaBC1 involved activation (phosphorylation in the Tyr 397) of Focal Adhesion Kinase (FAK) as one of the main adaptor proteins composing the mechanosensitive adhesome, which can directly regulate its catalytic activity through force-induced structural rearrangements^[37]^. In-Cell Western (ICW) results showed no significant effects on FAK phosphorylation after the simultaneous stimulation of the FN-binding integrins and NaBC1 transporter (Figure S2-e), suggesting that the B-induced mode of action of cell adhesion takes place via different pathways.

### 2.2 NaBC1 stimulation promotes myosin light chain phosphorylation

Force transmission after the assembly of focal adhesions is mediated by a Rho-associated kinase (ROCK) pathway, which in turn induces myosin light chain phosphorylation (pMLC) and increases cell contractility^[38,39]^. To determine whether the observed effect of active NaBC1 on cell adhesion had to do with tension, we analyzed cell cytoskeleton and myosin light chain phosphorylation. **Figure 2** shows non-transfected MSCs and transfected with esiRNA^NC^ and esiRNA^NaBC1^ respectively. PLLA-B2% and PLLA-B5% substrates presented significantly higher levels of pMLC and actin stress fibers than the PLLA and Glass control substrates (**Figure** 2-a,b and **S3**-a,b) in non-transfected and esiRNA^NC^ transfected cells. However, after transfecting cells with esiRNA^NaBC1^, pMLC levels on PLLA-B2% and PLLA-B5% substrates were reduced and resulted similar to PLLA and Glass, demonstrating that cells under NaBC1 stimulation respond by increasing intracellular tension and contractility.

**Figure 2.**
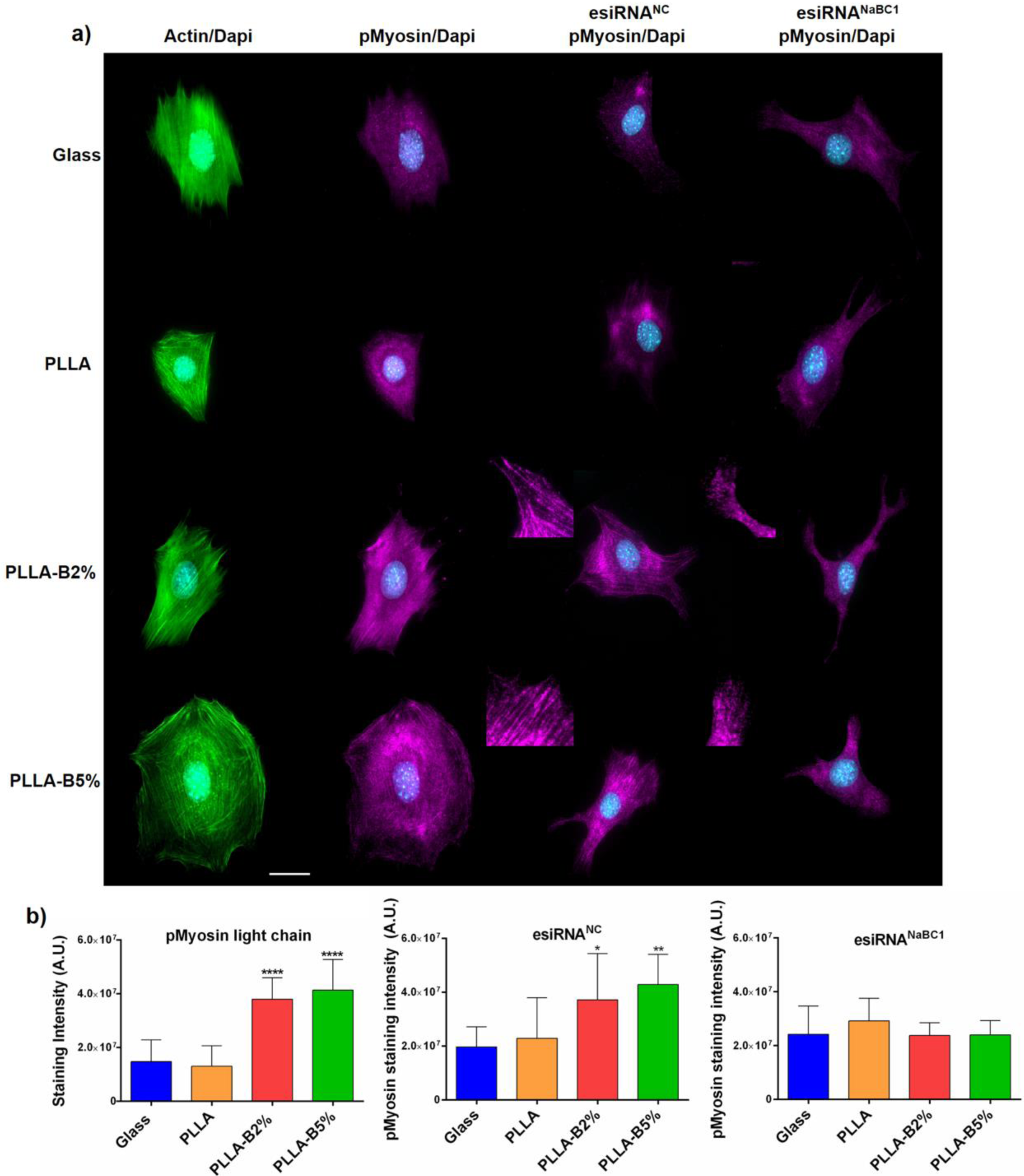
Boron effects on myosin light chain phosphorylation and actin stress fiber formation. a) Immunofluorescence images of actin cytoskeleton (green), nuclei (cyan) and pMLC (pMyosin, magenta) as markers of cellular tension and contractility. MSCs were cultured for 3 h onto functionalized substrates (FN-coated), with the presence of serum-depleted and boron (PLLA-B2%, PLLA-B5%) in the culture medium. The same experiment was performed after NaBC1 silencing using esiRNA^NC^ (negative control) and esiRNA^NaBC1^. The PLLA-B2% and PLLA-B5% substrates presented higher levels of actin fibers and pMLC staining that strongly diminished after NaBC1 silencing (see inset magnifications). Scale bar 25 µm. b) Image analysis quantification before and after NaBC1 silencing of pMLC staining, parameter related to cell contractility. Active NaBC1 induces cell contractility. Statistics are shown as mean ± standard deviation. n = 20 images/condition from three different biological replicas. Data was analyzed by an ordinary one-way ANOVA test and corrected for multiple comparisons using Dunnett analysis (P = 0.05). *p < 0.5, **p < 0.1, ****p < 0.0001.

To further investigate the role of active NaBC1 in cell contractility we worked with pharmacological inhibitors that impair contractility. We used Blebbistatin as specific inhibitor of myosin II activity (involved in cytokinesis and cell migration, cortical tension maintenance)^[40],^ and Y-27632 as specific inhibitor of Rho-kinase (disrupt myosin-dependent contractility)^[41]^. **Figure S4** shows immunofluorescence images of actin, vinculin and pMLC (pMyosin) after culturing MSCs with Blebbistatin (Figure S4-a) and Y-27632 (Figure S4-b). As expected, after contractility inhibition MSC morphology was affected in all conditions, however, vinculin and pMLC levels were significantly higher in PLLA-B2% and PLLA-B5% substrates compared with the reduced levels obtained in PLLA and Glass controls. Note that due to the lack of defined FA formation after using Blebbistatin and Y-27632, we could not perform their quantification. The fact that contractility inhibitors affect to a lesser extent the cells onto PLLA-B2% and PLLA-B5% substrates strongly supports the hypothesis that active NaBC1 reinforce cell adhesion and contractility at early cellular stages.

### 2.3. Active NaBC1 stimulates osteogenesis and inhibit adipogenesis in MSCs

We next evaluated the effect of simultaneous activation of FN-binding integrins and NaBC1 transporter on MSC differentiation. In order to explore the phenotypical and gene expression behavior of MSCs during commitment, we assayed three different experimental conditions: i) cells grown on basal medium (hereafter Basal), composed of a conventional growing medium containing 10% FBS without any supplements or growth factors; ii) cells grown on an osteogenic-specific differentiation medium (hereafter Ob); iii) cells grown on an adipogenic-specific differentiation medium (hereafter Ad). For the basal conditions and to promote osteogenesis, cells were seeded at 10,000 cells cm^-2^, whereas cells were seeded at 30,000 cells cm^-2^ to favor adipogenesis^[42]^. **Figure S5**-a shows immunofluorescence images of MSCs cultured for 3 days onto FN-coated substrates and (with/without) boron under Basal and Ob conditions for evaluation of Runt-related transcription factor 2 (Runx2) as an early expressed transcription factor in osteogenenesis^[43]^. The presence of boron did not increase Runx2 expression either under Basal conditions or under osteogenic stimulation (Ob).

Figure S5-b shows the qPCR analysis of specific gene encoding transcription factors involved in the early onset of lineage commitment versus osteogenic (Runx2) and adipogenic (adipocyte peroxisome proliferation-activated receptor-PPARγ2)^[44]^ lineages, under Basal and Ob or Ad conditions. In both cases, boron did not increase either Runx2 or PPARγ2 expression alone, or after the induction of lineage commitment, suggesting that NaBC1 stimulation has no differential effect on the gene expression of these transcription factors at this experimental time point. Previous reports have contrasted the general assumption of the role of Runx2 levels in mediating osteogenesis, showing Runx2 constitutive expression and no correlation between Runx2 mRNA/protein levels and the osteoblast phenotype^[43]^.

**Figure 3**-a shows representative images of late expressed markers involved in osteogenic (osteopontin-OPN and alkaline phosphatase-ALP)^[45]^ and adipogenic (adipocyte formation) differentiation after 15 days of culture. MSCs were cultured for 15 days onto FN-functionalized substrates and boron under Basal and Ob or Ad conditions. NaBC1 stimulation induced higher OPN and ALP staining levels in an osteogenic defined medium than in PLLA and Glass control conditions. Surprisingly, the capacity of the sole presence of boron to promote osteogenic lineage commitment was particularly strong, as revealed by the higher levels of OPN and ALP under Basal conditions (Figure 3-a). We have obtained similar results after histological analysis of Alizarin red, ALP and Von Kossa staining (Figure S5-c) and immunofluorescence detection of Collagen I (Col I), Integrin Binding Sialoprotein (IBSP) and Osteocalcin (OCN) (Figure S5-d). Conversely, MSCs cultured for 15 days onto FN-functionalized substrates and boron under Ad conditions gave the opposite results. Adipocyte formation after Oil Red O staining was evident only in the Glass and PLLA substrates, showing the clear formation of rounded cells with lipid vacuoles containing fat droplets^[46]^. PLLA-B2% and PLLA-B5% substrates inhibited adipocyte formation, and kept the polygonal morphology typical of osteoblasts with a minimal amount of red-stained fat droplets (Figure 3-a).

**Figure 3.**
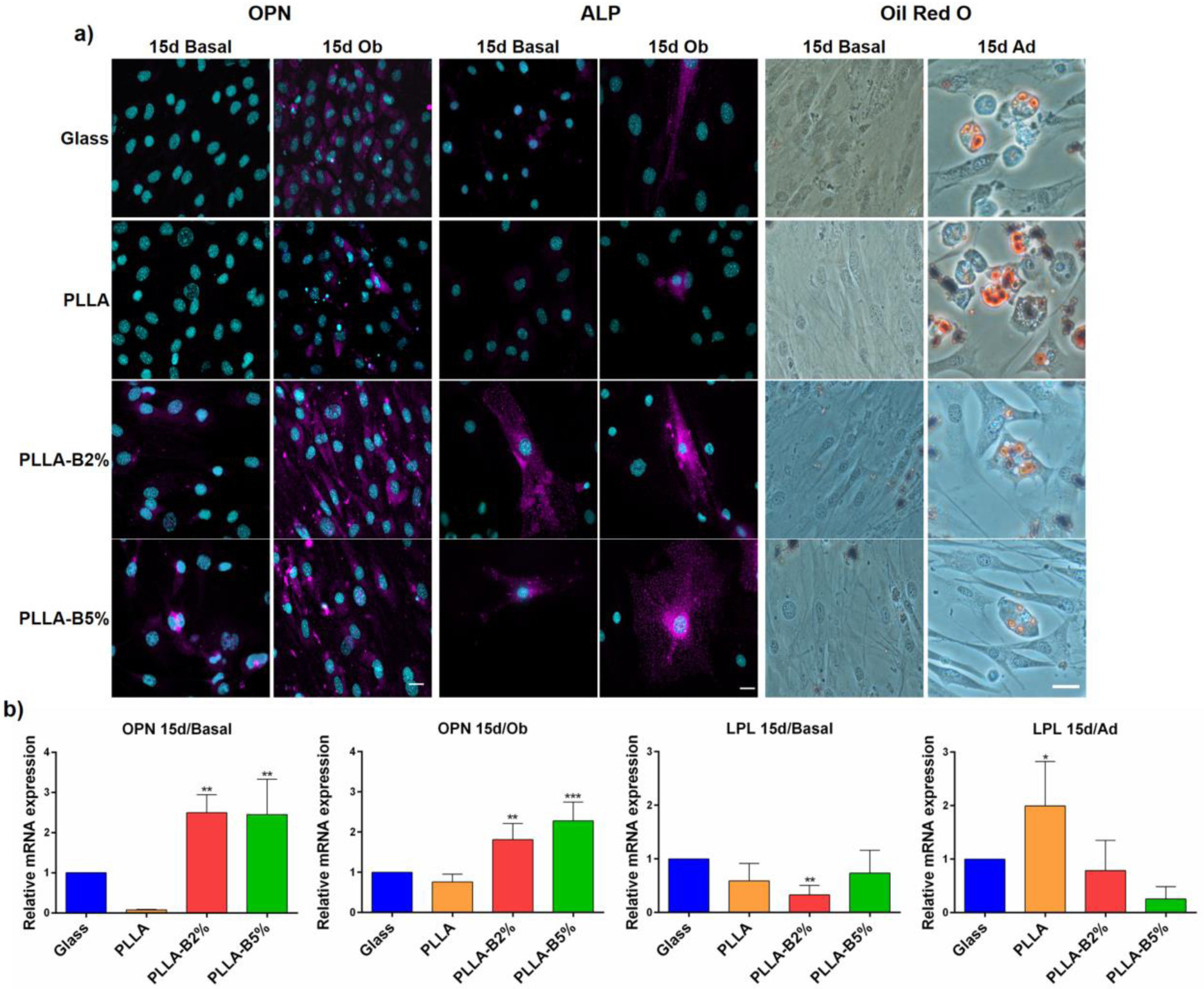
Boron effect on MSC differentiation. a) Immunofluorescence images of MSCs cultured for 15 days onto functionalized substrates (FN-coated) and boron (PLLA-B2%, PLLA-B5%) in culture medium, under basal and differentiation conditions (osteogenic), for late osteogenic detection markers (OPN and ALP-magenta, nuclei-cyan). Scale bar 25 µm. Oil red O staining of MSCs cultured for 15 days onto functionalized substrates (FN-coated) and boron (PLLA-B2%, PLLA-B5%) in culture medium, under basal and differentiation conditions (adipogenic), for adipocyte detection showing typical rounded adipocyte shape and accumulation of intracellular fat droplets. Scale bar 25 µm. b) qPCR analysis of relative mRNA expression of late expressed markers involved in osteogenic (OPN) and adipogenic (LPL) lineage determination. RNA was extracted after 15 days of culture under both basal and differentiation conditions (osteogenic and adipogenic differentiation media). Statistics are shown as mean ± standard deviation. n = 5 different biological replicas. Data was analyzed by an ordinary one-way ANOVA test and corrected for multiple comparisons using Tukey analysis (P = 0.05). *p < 0.05, **p < 0.01, ***p < 0.001.

We also investigated the gene expression of the late expressed markers involved in osteogenic (osteopontin-OPN) and adipogenic (Lipoprotein lipase-LPL)^[47]^ differentiation by qPCR after 15 days of culture. Figure 3-b shows that NaBC1 and FN-binding integrin co-stimulation strongly induced OPN expression under Basal and osteogenic differentiation conditions, whereas they extremely reduced LPL levels, results that are in concordance with the immunofluorescence images shown in Figure 3-a and those obtained for focal adhesion formation.

These results demonstrate that the simultaneous stimulation of NaBC1 and FN-binding integrins promotes osteogenesis (in basal conditions) and strongly inhibits adipogenesis, even in an adipogenic induction medium.

### 2.4 Boron up-regulates NaBC1 and FN-binding integrins expression in MSC

The sodium-boron co-transporter NaBC1, essential for boron homeostasis, has already been characterized^[18]^. We recently reported a 3D molecular model based on the sequence homology of other bicarbonate transporter-related proteins^[22]^. The NaBC1 transporter is boron-specific and functions as an obligated Na^+^-coupled borate cotransporter. As we used Sodium Tetraborate Decahydrate (borax) in this work, we confirmed that borax is the natural ligand for NaBC1 activation and that its transport is not through other mechanisms, such as membrane passive diffusion, as with boric acid^[18]^.

Although there is some evidence for the tissue ubiquity of NaBC1^[47]^, we confirmed its presence in our cellular system under the same experimental conditions (after 3 or 15 days of culture and under basal or differentiation media) used in previous studies. When MSCs NaBC1 mRNA were amplified by qPCR, the results showed that in all the conditions assessed, NaBC1 gene expression was upregulated exclusively by boron and that this upregulation was dose-dependent (**Figure S6**-a). We have also confirmed NaBC1 upregulation at protein level by In-Cell Western (Figure S6-b).

We next explored whether boron activation of NaBC1 influences the expression of FN-binding integrins. Previous studies have reported the critical role of α_5_β^[48]^ and α_v_β_3_^[49]^ integrins in osteogenic-adipogenic balance, as well as in regulating gene expression by ion-channels, phenomenon only described for potassium channels^[50,51]^. **Figure 4**-a shows immunofluorescence images and staining intensity of α5 and αv integrins and reveals enhanced integrin levels for PLLA-B2% and PLLA-B5% substrates. We have also detected α_5_ and α_v_ integrins at protein level in PLLA-B2% and PLLA-B5% substrates by In-Cell Western (**Figure S7**-a).

**Figure 4.**
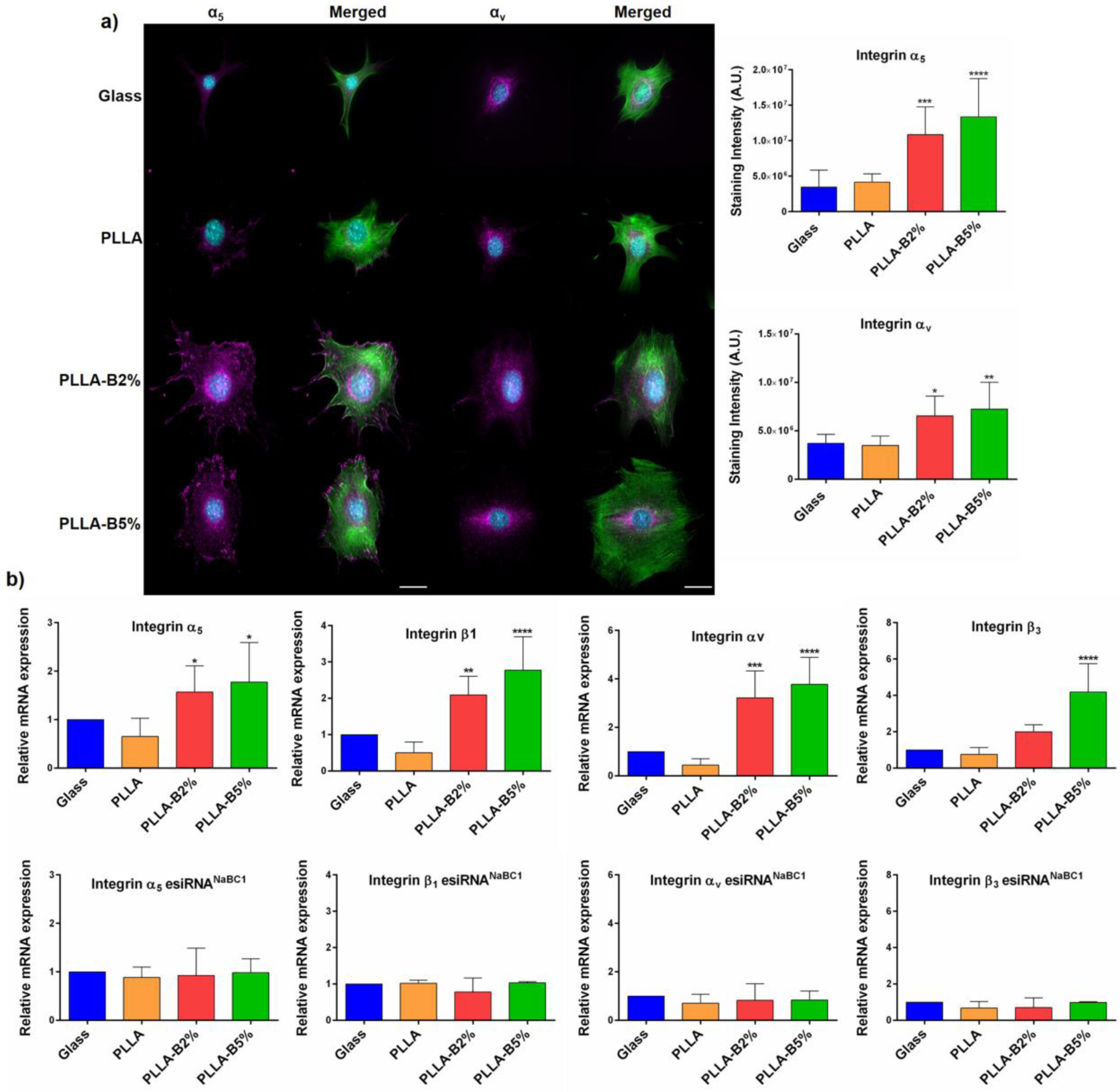
NaBC1 induces FN-binding integrins expression. a) Immunofluorescence images of MSCs cells cultured for 3 h onto functionalized substrates (FN-coated), serum-depleted and boron (PLLA-B2%, PLLA-B5%) in culture medium. Images show actin cytoskeleton (green) and α_5_ or α_v_ integrins (magenta). Boron induces the expression of FN-binding integrins. Scale bar 25 µm. Right graphs: image analysis quantification of α_5_ and α_v_ integrins levels. n = 20 images/condition from 3 different biological replicas. b) qPCR analysis of relative mRNA expression of FN-binding integrins (α_5_β_1_ and α_v_ β_3_) before and after silencing of NaBC1 transporter. Active NaBC1 induces FN-binding integrins expression. n = 6 different biological replicas. Statistics are shown as mean ± standard deviation. Data was analyzed by an ordinary one-way ANOVA test and corrected for multiple comparisons using Tukey analysis (P = 0.05). *p < 0.05, **p < 0.01, ***p < 0.001, ****p < 0.001.

We further analyzed mRNAs of FN-binding integrins from non-transfected MSCs and transfected with esiRNA^NC^ and esiRNA^NaBC1^ respectively. qPCR analysis showed that active NaBC1 induced gene expression levels of α_5_β_1_ and α_v_β_3_ integrins (Figure 4-b), effect that was abrogated after NaBC1 silencing. This fact demonstrated that NaBC1 activation is capable of upregulate integrin gene expression as previously reported for other ion-channels^[50]^.

Collectively, these findings demonstrate that boron stimulation of NaBC1 leads to enhanced expression of α_5_β_1_ and α_v_β_3_ integrins, consistent with enhanced osteogenesis (Figure 3), larger spreading area and the greater number of large focal adhesions formed (Figure 1) and increased intracellular tension (Figure 2).

### 2.5. Active NaBC1 co-localizes with FN-binding integrins and BMPR1A receptors

We next wanted to study the interplay between NaBC1 and the other membrane receptors involved in the osteogenic pathway. To test whether the simultaneous stimulation of NaBC1 and the α_5_β_1_ and α_v_β_3_ integrins followed a physical protein-protein interaction, as has previously been described for other ion-channels^[52]^, we performed NaBC1/α_5-_α_v_ co-localization assays using the DUOLINK® PLA kit system, which shows only the positive signals generated between two different proteins less than 40 nm apart. Similarly, we also investigated NaBC1 co-localization with BMPR1A as the main membrane growth factor receptor that it has been shown to co-localize with integrin α_v_ to induce osteogenesis, even in absence of its BMP ligands^[8]^ members of the transforming growth factor β (TGFβ) superfamily^[53]^.

**Figure 5**-a shows the co-localization of NaBC1 with α_5_ and α_v_ integrins as well as co-localization of NaBC1 with BMPR1A. A few fluorescent dots were present in the Glass and PLLA substrates, however, the positive signals in the PLLA-B2% and PLLA-B5% samples increased in a dose-dependent manner, indicating that NaBC1 stimulation promotes the effective co-localization of the receptors, considering that FN-binding integrins are also enhanced by the presence of B on the FN-coated samples. Surprisingly, NaBC1/BMPR1A co-localization occurred even without external addition of BMPs, the BMPR1A natural ligand, as previously reported for osteogenic nanotopographical surfaces^[8]^. We have also detected an increase of α_5_ and α_v_ integrins and BMPR1A at protein level in PLLA-B2% and PLLA-B5% substrates by In-Cell Western (Figure S7-b). It should be noted that BMPs were not used in this experiment as supplemented-growth factors to induce osteogenesis. These results describe for the first time a new protein-protein interaction between NaBC1/BMPR1A as an osteogenic induction mechanism, suggesting that NaBC1 acts as a novel mechanoreceptor that activates BMPR1A in crosstalk with FN-binding integrins, and in the absence of external growth factors.

**Figure 5.**
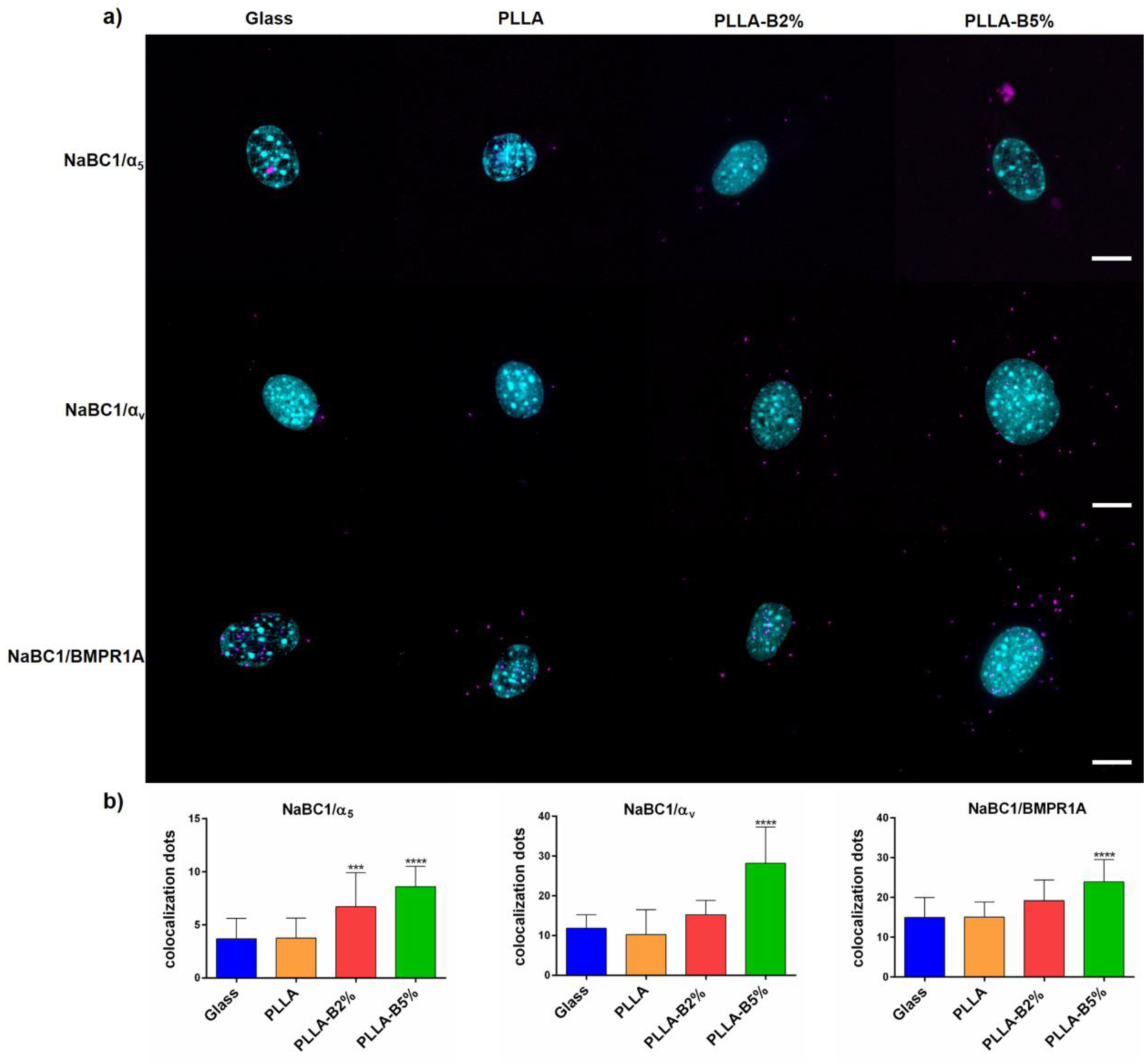
NaBC1/α_5,_ α_v_ and NaBC1/BMPR1A co-localization. a) Images of nuclei (cyan) and co-localization dots (magenta) of MSCs cultured for 3 h onto functionalized substrates (FN-coated), serum-depleted and boron (PLLA-B2%, PLLA-B5%) in the culture medium. Scale bar represents 10 µm. b) Image analysis quantification of the number of co-localization dots of NaBC1/α_5_, NaBC1/α_v_ and NaBC1/BMPR1A. Co-localization levels increase with boron concentration. n = 30 images/condition from 3 different biological replicas. Statistics are shown as mean ± standard deviation. Data was analyzed by an ordinary one-way ANOVA test and corrected for multiple comparisons using Tukey analysis (P = 0.05). ***p < 0.001, ****p < 0.0001.

### 2.6. Active NaBC1 induced ERK1/2-Smad1-Akt phosphorylation and active YAP-Smad1 nucleus translocation

We next wanted to understand the downstream signaling that was triggered by NaBC1 activation leading to osteogenesis. For this, we first explored the phosphorylation of the main effector molecules involved in osteogenic commitment, such as Runx2, Extracellular signal-Regulated Kinase 1/2 (ERK1/2), Protein kinase B (Akt) and Small Mothers Against Decapentaplegic 1 (Smad1/5/8). In order to confirm that boron activation of NaBC1 relies on intracellular tension, we also evaluated the active Yes Associated Protein 1 (YAP) presence as another force-induced mechanotransductive marker. In-Cell Western experiments showed that NaBC1 activation significantly induced ERK1/2, Akt, Smad1 phosphorylation (**Figure 6**-a) and the levels of active YAP, while it had no effect on Runx2.

**Figure 6.**
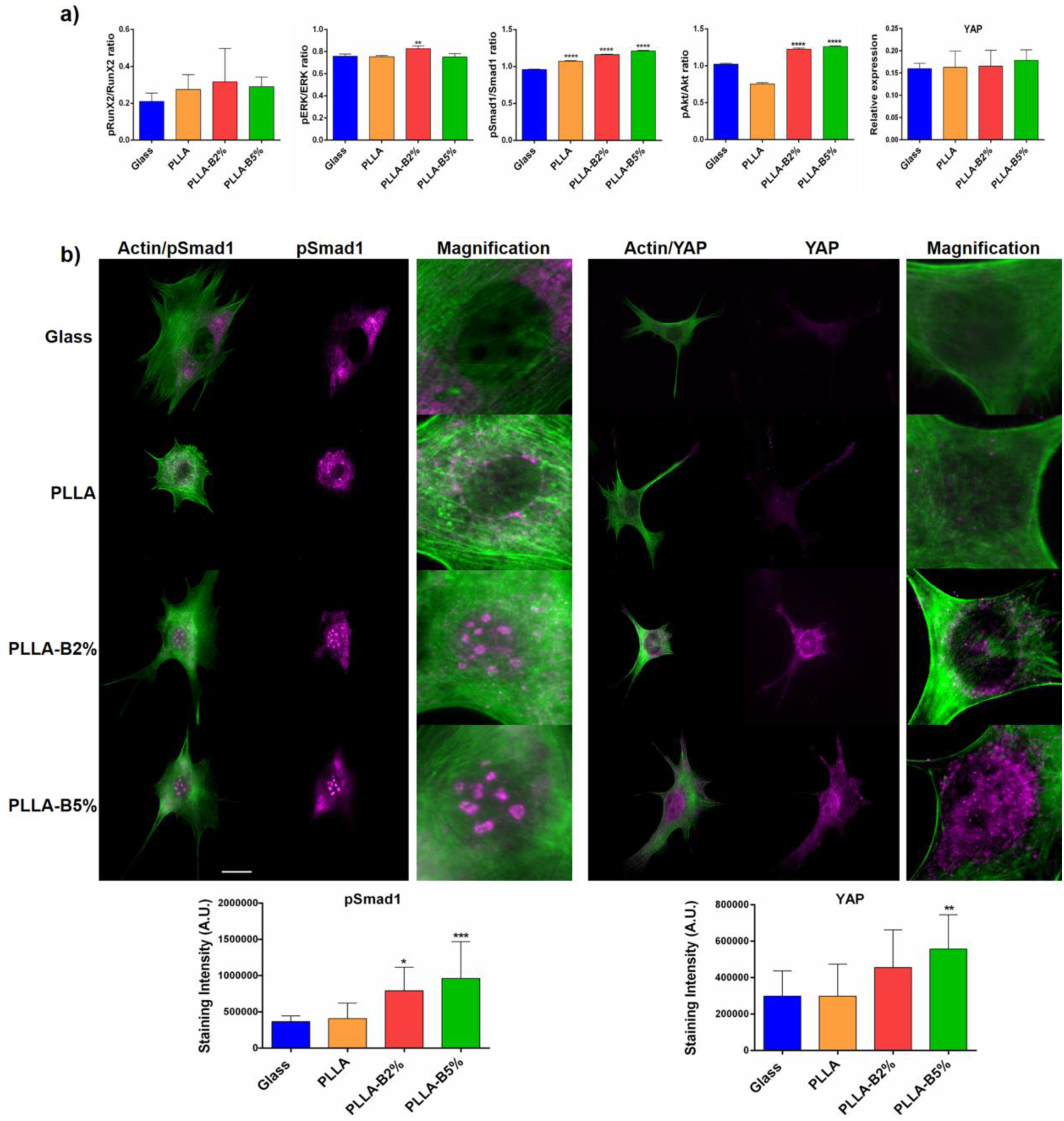
Effects of NaBC1 activation in intracellular signaling. a) In-Cell Western assay showing pRunx2/Runx2, pERK/ERK, pAkt/Akt and pSmad1/Smad1 ratios and active form of YAP on MSCs cultured onto functionalized substrates (FN-coated) serum-depleted and boron (PLLA-B2%, PLLA-B5%) in the culture medium after 90 minutes of culture. Simultaneous stimulation of FN-binding integrins and NaBC1 resulted in a significant enhancement of ERK, Smad1 and Akt phosphorylation as well as active YAP detection. Statistics are shown as mean ± standard deviation. n = 4 biological replicas. Data was analyzed by an ordinary one-way ANOVA test and corrected for multiple comparisons using Tukey analysis (P = 0.05). **p < 0.01, ****p < 0.0001. b) Immunofluorescence images of MSCs cells cultured for 3 h onto functionalized substrates (FN-coated), serum-depleted and boron (PLLA-B2%, PLLA-B5%) in culture medium. Images show actin cytoskeleton (green), pSmad1 and YAP (magenta). Boron induces active Smad1 and YAP translocation into the cell nucleus (see inset magnifications). Scale bar 25 µm. Image analysis quantification of pSmad1 and active YAP levels. Statistics are shown as mean ± standard deviation. n = 20 images/condition from 3 different biological replicas. Data was analyzed by an ordinary one-way ANOVA test and corrected for multiple comparisons using Tukey analysis (P = 0.05). *p < 0.05, **p < 0.01, ***p < 0.001.

We further evaluated active Smad1 and YAP intracellular location, as Smad1 tethers with other Smad4 proteins and translocate to the nucleus after phosphorylation, becoming transcriptionally active for the expression of genes involved in osteogenic commitment^[54]^, and YAP is a transcriptional co-activator acting downstream of the Hippo pathway and one of its substrates is the active nuclear pSmad1^[55]^. It has been described that high mechanical stress promotes nuclear localization of YAP to drive osteogenesis^[56]^. Figure 6-b shows clearly active Smad1 and YAP nuclear accumulation with a considerable increase only in PLLA-B2% and PLLA-B5% substrates (see magnification images right panel).

## 3. Discussion

Understanding the role of ECM and its ability to interact with membrane receptors to mimic natural cellular niches is critical for the success of tissue engineering approaches. Diverse strategies have been used to control MSC growth and differentiation, including: modulation of the physical properties of materials in terms of chemistry^[4]^, stiffness^[5,6]^ and topography^[7,8]^, controlling spatio-temporal GF delivery/exposure to mimic cell receptor-GF crosstalk and activate the signaling pathways involved in MSC fate^[9]^, or even combining modifications of their material properties by soluble GF approaches^[57]^. As has been broadly described in the literature, cell mechanotransduction mechanisms are the key determinants of MSC physiology^[5,11,57,58]^. Even though several reports have described ion-channels as mechanosensors^[15,16]^, whether their contribution to MSC behavior is caused by force transmission or is simply due to the role of ion-homeostasis in intracellular mechanisms is still not clear, and the studies published to date focus mainly on Ca^2+^, K^+^ and Cl^-^ channels^[17]^.

The present study demonstrates that the boron-channel, a NaBC1 transporter, acts by co-localizing with BMPR1A and activating the BMP-dependent downstream pathways in the absence of any external BMP supplementation, driving the MSCs to osteogenic commitment. As BMP2 and 4 are known to be constitutively synthesized by C3H10T1/2 in a given osteogenic medium^[59]^, the role of BMPs in BMPR1A activation in long term cultures for MSC differentiation under certain conditions cannot be ruled out. However, as in our co-localization and In-Cell Western experiments we used serum-depleted basal media only, without any osteogenic induction, we can propose a new crosstalk mechanism involving active NaBC1 and BMPR1A in the absence of external BMPs.

Recent reports have described the effects of boron in the induction of osteogenesis^[23-25]^ and inhibition of adipogenesis^[26,27]^. Even though they do describe the possible metabolic pathways that explain the observed effects (mainly on adipogenic inhibition), they do not describe the mechanism used to transport boron and the intracellular signaling mechanisms induced after activation of NaBC1 transporter.

Our initial hypothesis was that the simultaneous activation of membrane receptors enhances intracellular signaling, as previously described for cooperation mechanisms between integrins and growth factor receptors^[60]^. To test this hypothesis, we employed FN-coated substrates for α_5_β_1_ and α_v_β_3_, and boron for NaBC1 stimulation, respectively. In the current study, we used borax solution (Na_2_B_4_O_7_), so that the B uptake by the cells would be guaranteed via NaBC1, since this is an obligated Na^+^-coupled borate co-transporter^[18]^. We have shown that the simultaneous activation of NaBC1 and FN-binding integrins enhances cell adhesion in terms of greater cell spreading and the formation of mature focal adhesions (Figure 1 and S2). The transient depletion of NaBC1 function, after transfecting cells with esiRNA^NaBC1^, clearly points out the synergistic effect of NaBC1 transporter in combination with FN-binding integrins, and strongly suggest that NaBC1 activation regulates cell adhesion. This enhanced adhesion (in FA size and number) determines the MSCs differentiation towards osteogenic commitment, while it inhibits adipogenesis, as expected, being the osteogenic-adipogenic balance is mutually exclusive (Figure 3).

Our results fit in nicely with previous reports describing the importance of adhesion size, normally associated with cell motility (small adhesions, low intracellular tension derives into adipocytes) or high stability and cytoskeletal tension (long and mature adhesions, high intracellular tension derives into osteoblasts) ^[58]^. Despite our findings on adhesion, we did not find any significant differences in the pFAK/FAK ratio (Figure S2-e). As FAK is one of the main downstream effectors after integrin activation^[7]^, this result was unexpected, and suggests that simultaneous stimulation of NaBC1 and FN-binding integrins involves other metabolic pathways rather than FAK phosphorylation, as we previously reported^[21]^. To verify this finding we explored the phosphorylation of the myosin light chain (pMLC) involved in cell contractility and acting downstream of the FAK. The results show that NaBC1 activation induced elevated levels of pMLC and actin stress fibers (Figure 2), effects reverted after NaBC1 silencing, specifying NaBC1 activation in the promotion of cell adhesion and contractility. The fact that after using contractility inhibitors pMLC and vinculin levels remained elevated when simultaneous stimulation of NaBC1 and FN-binding integrins occurred, strongly suggest that the formation of integrin-cytoskeleton linkages is immediate in PLLA-B2% and PLLA-B5% substrates. Cooperation between NaBC1/FN-binding integrins can produce a more robust strengthening response.

Our results show that the simultaneous activation of NaBC1 and FN-binding integrins causes an adhesion-primed state of the MSCs related to intracellular tension, caused by the increased FA, pMLC and stress actin fibers, and elevated α_5_β_1_/α_v_β_3_ integrins and NaBC1 expression (Figure 4 and S6). NaBC1/α_5_β_1_/α_v_β_3_ and NaBC1/BMPR1A clustering and co-localization (Figure 5) may act as a mechanosensing switch affecting the downstream effectors of TGFβ and Hippo pathways, Smad1 and YAP respectively, which became active and translocate to the nucleus to exert their transcriptional function^[54,55]^ (Figure 6). However, further studies will be needed to establish the precise role of NaBC1 as a mechanosensor, including their response on surfaces of controlled elastic properties.

Some studies have described the role of FN-binding integrins in osteogenic-adipogenic balance^[48,61,62]^, the role of BMPR1A driving osteogenesis after BMP ligand activation^[63]^, and describing YAP as the key determinant of cell mechanics that controls focal adhesion assembly^[64]^. Even though Runx2 is a key target for active Smad1 and has been proposed as the main mediator of downstream BMP actions, several authors report that some BMP effects are Runx2-independent in bone formation^[59]^. We did not find significantly high levels of this osteogenic regulator gene in our experimental system, but instead we did find higher OPN levels expression (even in Basal conditions). OPN-integrin interaction has been described as critical for MSCs osteogenic differentiation^[49]^, and thus higher OPN expression, obtained by the sole addition of boron (Figure 3) supports the crosstalk mechanism between active NaBC1/FN-binding integrins/BMPR1A. Another pSmad1 target is Hoxc-8, a transcriptional repressor that liberates the transcription of the OPN gene^[65]^ after pSmad1 binding and agrees with our hypothesis.

We have therefore demonstrated a novel function for the NaBC1 ion-channel in MSCs. To date, the synergistic interactions between ion-channels and other membrane receptors have not been exploited to engineer material systems for biomedical applications. We here propose a simple approach for driving osteogenesis (and in turn inhibiting adipogenesis) through a mechanism that involves the simultaneous stimulation of NaBC1, BMPR1A and α_5_β_1_/α_v_β_3_ integrins to enhance intracellular signaling. As the co-localization experiments show that these receptors interact physically and activates the translocation of Smad1 and YAP into the nucleus, we thus propose NaBC1 as a novel mechanosensor. Our results indicate an adhesion-primed state related to intracellular tension involving the diverse crosstalk mechanisms described in **Figure 7**. These findings will open up new ways for engineering biomaterials to be employed in biomedical applications avoiding the use of high doses of soluble GFs.

**Figure 7.**
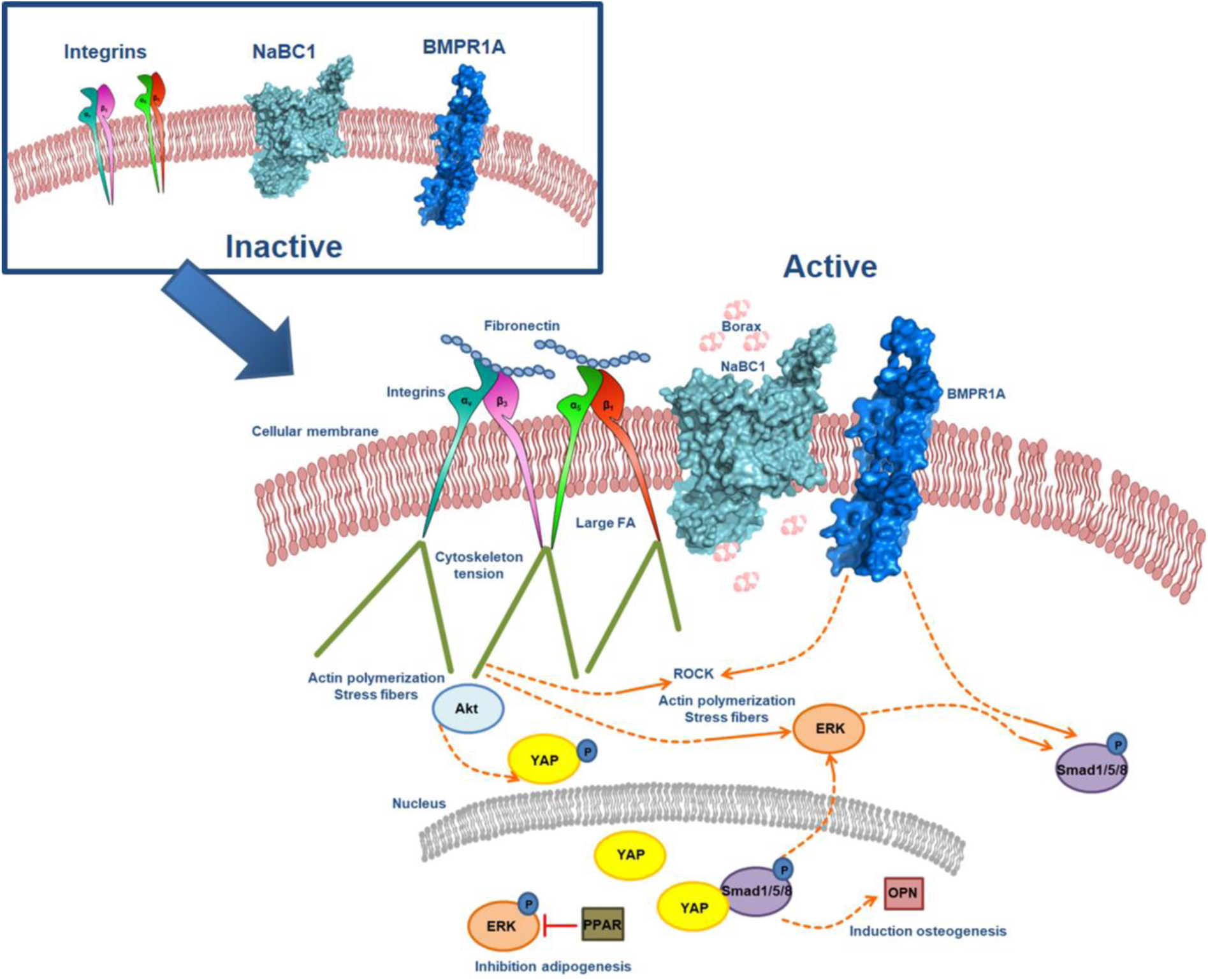
Molecular control of spatiotemporal cues for synergistic intracellular signaling. Description of molecular events occurring after simultaneous NaBC1/α_5_β_1_/α_v_β_3_ and NaBC1/BMPR1A activation. FN stimulate FN-binding integrins while boron stimulates the NaBC1 transporter. NaBC1 co-localizes with FN-binding integrins and BMPR1A, thus stimulating, after physical interaction, downstream pathways controlling MSCs fate. Canonical BMPR1A activation concludes in Smad1 phosphorylation, inducing osteogenesis via OPN expression. After ligand binding (FN) and NaBC1 co-localization, active integrins cluster and form large focal adhesions, producing high intracellular tension that derive into YAP translocation to the nucleus and induction of osteogenesis. High mechanical tension activates ERK, which inhibits adipogenesis after phosphorylation.

## 4. Conclusions

This work proposes a novel way to control MSC fate using the interplay between specific cell membrane receptors to control the MSC fate. Our data show that the simultaneous stimulation of NaBC1 and FN-binding integrins generates an adhesion-primed state (mature FA, cell spreading), stimulating the mechanical determinants that affect cell fate through actomyosin forces (pMLC, actin stress fibers), and the mechanosensitive translocation of transcription factors (pSmad, YAP). This stimulation enhances osteogenesis and inhibits adipogenesis in the absence of other soluble chemicals or GFs. We describe a novel mechanism involving the crosstalk and co-localization of active NaBC1/α_5_β_1_/α_v_β_3_ integrins and NaBC1/BMPR1A and the intracellular pathways involved, with a novel function for NaBC1 as a mechanosensitive ion-channel.

## 5. Experimental Section

### 5.1. Material substrates

Cleaned glass cover slips were used as control substrates. PLLA (Cargill Dow) was dissolved at a concentration of 2% (w/v) in chloroform. Sodium Tetraborate Decahydrate Borax 10 Mol (hereafter boron) (Na_2_B_4_O_7_·10H_2_O, Borax España S.A) was dissolved in the culture medium in all the experiments. Two different boron concentrations were used, 0.59 mM and 1.47 mM respectively, in the PLLA-B2% and PLLA-B5% samples. PLLA 2% (w/v) solution was used to prepare thin films by spin-coating on cleaned glass cover slips for 30 s at 3000 rpm. Samples were dried at 60 °C *in vacuo* for 4 h.

All the substrates were functionalized with human plasma fibronectin (FN, Sigma-Aldrich). After sterilizing with UV for 30 min, the substrates were coated with 20 µg mL^-1^ FN solution in Dulbecco’s Phosphate Saline Buffer (DPBS) for 1 h at room temperature.

### 5.2. Cytotoxicity assay

MTS quantitative assay (The CellTiter 96 Aqueous One Solution Cell Proliferation Assay, Promega) was performed to assess cytocompatibility of boron with MSCs and to establish the maximum working concentrations to use with this particular cell line. 20,000 cells cm^-2^ were seeded onto a p-24 multi-well plate and metabolic activity was measured after 24 h of incubation of cells with different quantities of boron in media (0.2, 0.6, 3, 6, 10 and 20 mg mL^-1^). Cells were then incubated for 3 h with MTS (tetrazolium salt) at 37 °C and the formation of formazan was followed by measuring absorbance at 490 nm. All measurements were performed in triplicate.

### 5.3. MSCs culture

Murine embryonic mesenchymal stem cells (MSCs) C3H10T1/2 (RIKEN Cell Bank, Japan) were cultured in Dulbecco’s Modified Eagle Medium (DMEM) with high glucose content, 10% fetal bovine serum (FBS) and 1% antibiotics (penicillin/streptomycin) at 37 °C in a humidified atmosphere of 5 % CO_2_ (conventional growth medium and basal medium composition (hereinafter Basal). Cells were subcultured once a week before reaching confluence.

### 5.4. MSC transfection

MSCs were seeded at 60,000 cells cm^-2^ in Dulbecco’s Modified Eagle Medium (DMEM) with high glucose content, 10% fetal bovine serum (FBS) and 1% antibiotics (penicillin/streptomycin) at 37 °C in a humidified atmosphere of 5 % CO_2_. After 24 h, cells were washed with Opti-MEM reduced serum medium (ThermoFisher) and transfected using pre-designed MISSION esiRNA (Sigma-Aldrich) against mouse NaBC1 transporter in X-tremeGENE siRNA Transfection Reagent (Roche), following manufacturer’s instructions. MISSION siRNA Fluorescent Universal Negative Control 1, Cyanine 3 (NC, Sigma-Aldrich) was used as a control of transfection efficiency.

### 5.5. Cell adhesion experiments

For cell adhesion experiments, MSCs or esiRNA transfected MSCs were plated on glass covers and PLLA substrates functionalized with FN at a density of 5,000 cells cm^-2^. Cells were cultured in DMEM with high glucose content, 1% antibiotics and in absence of serum. Samples were also supplemented with 0.59 mM or 1.47 mM from borax solution as required. After 3 h at 37 °C cells were fixed and immunostained.

### 5.5. Cell differentiation experiments

For differentiation experiments, MSCs were plated on glass covers and PLLA substrates functionalized with FN at a density of 20,000 cells cm^-2^ for osteogenic and basal conditions and 30,000 cells cm^-2^ for adipogenic differentiation. Cells were cultured for 48 h in DMEM growth media until they reached 70–80 % confluence. After confluence, differentiation was induced with osteogenic media (hereinafter Ob) (DMEM growth medium supplemented with Ascorbic Acid 50 µg mL^-1^, Glycerophosphate 10 mM and Dexamethasone 0.1 µM) or adipogenic media (hereinafter Ad) (DMEM growth medium supplemented with 3-isobutyl-1-methyl-xanthine (IBMX) 0.5 mM, indomethacin 60 µM and Hydrocortisone 0.5 µM). Samples were also supplemented with 0.59 mM or 1.47 mM of borax solution in every change of medium as required. Media was changed every 3 days until end-point assay. All differentiation experiments were finished after 3 days for analysis of early transcription factors (Runt-related transcription factor 2 - Runx2 - and Peroxisome proliferator-activated receptor gamma - PPAR-γ) or 15 days for analysis of osteogenic/adipogenic markers (Osteopontin – OPN, Osteocalcin-OCN, Integrin Binding Sialoprotein-IBSP, Alkaline Phosphatase-ALP, Lipoprotein lipase - LPL or adipocyte formation).

### 5.4. Immunohistochemistry assays and staining

After culture, cells were fixed in a 4% formalin solution (Sigma-Aldrich) at 4 °C for 30 min. Samples were then rinsed with DPBS and permeabilized with DPBS/0.5% Triton X-100 at room temperature for 5 min. Samples were then incubated with primary antibodies in blocking buffer DPBS/2% BSA at 37 °C for 1 h or overnight at 4 °C. The samples were then rinsed twice in DPBS/0.1% Triton X-100 and incubated with the secondary antibody and/or BODIPY FL phallacidin (Invitrogen, 1:100) at room temperature for 1 h. Finally, samples were washed twice in DPBS/0.1% Triton X-100 before mounting with Vectashield containing DAPI (Vector Laboratories) and observed under an epifluorescence microscope (Nikon Eclipse 80i).

For cell adhesion studies monoclonal antibodies against vinculin-FA detection (Sigma-Aldrich, 1:400), pMLC-intracellular tension (Cell Signaling, 1:200), integrin α_5_ (abcam, 1:500), integrin α_v_ (abcam, 1:500), pSmad1 (Cell Signalling, 1:200), active YAP (abcam, 1:500) were used. Cy^3^ conjugated (Jackson Immunoresearch, 1:200) or Alexa Fluor 555 (ThermoFisher, 1:700) were used as a secondary antibodies.

Several specific markers were used to evaluate differentiation. Osteogenic differentiation was assessed by Runx2 (abcam, 1:100), Osteopontin (Santa cruz Biotechnology, 1:100), Osteocalcin (abcam, 1:200), Integrin Binding Sialoprotein (Santa Cruz Biotechnology, 1:200), Collagen I (abcam, 1:200) as primary antibodies. Cy^3^ (Jackson Immunoresearch, 1:200) and Alexa Fluor 488 or 555 (Invitrogen, 1:700) were used as secondary antibodies.

Osteogenic differentiation was assessed also by histological staining.

Alkaline phosphatase activity (ALP) was determined staining samples with naphtol AS-MX/Fast Red TR (Sigma-Aldrich) for 30 minutes at room temperature. After incubation, samples were washed and nuclei were labelled with Hoechst. These samples can be visualized either histologically or by fluorescence microscopy.

Von Kossa and Alizarin red staining were performed following the PROMOCELL procedure. Adipogenic differentiation was detected by observation of lipid levels that were qualitatively assessed by a standard Oil Red O staining protocol. Briefly, cells were washed in DPBS. Immediately before use, 30 mL of a stock solution of Oil Red O (3 mg mL^-1^ in 99 % isopropanol) was mixed with 20 mL diH_2_O, filtered and applied for 10-15 min at room temperature to cells pre-equilibrated with 60% isopropanol.

### 5.5. Gene expression

Total RNA was extracted from MSCs cultured for 3 or 15 days under different experimental conditions using RNeasy Micro Kit (Qiagen). RNA quantity and integrity was measured with a NanoDrop 1000 (ThermoScientific). Then 500 ng of RNA were reverse transcribed using the Superscript III reverse transcriptase (Invitrogen) and oligo dT primer (Invitrogen). Real-time qPCR was performed using Sybr select master mix and 7500 Real Time PCR system from Applied Biosystems. The reactions were run at least in triplicate for both technical and biological replicas. The primers used for amplification were designed from sequences found in the GenBank database and included:

Runx2 (NM_001146038.1, Forward: 5’-TGA GAG TAG GTG TCC CGC CT-3’, Reverse: 5’-TGT GGA TTA AAA GGA CTT GGT GC-3’) and Osteopontin (NM_001204201.1, Forward: 5’-TTT GCC TGT TTG GCA TTG C-3’, Reverse: 5’-TGG GTG CAG GCT GTA AAG CT-3’) for osteogenic differentiation. PPARγ2 (NM_001127330.1, Forward: 5’-AGC AAA GAG GTG GCC ATC C-3’, Reverse: 5’-CTT GCA CGG CTT CTA CG-3’) and LPL (NM_008509.2, Forward: 5’-TGC CCT AAG GAC CCC TGA A-3’, Reverse: 5’-CAG TTA GAC ACA GAG TCT GC-3’) were used for adipogenic differentiation.

NaBC1 (NM_001081162.1, Forward: 5’-GAG GTT CGC TTT GTC ATC CTG G-3’, Reverse: 5’-TTC CTC TGT GCG AGT CTT CAG G-3 were used for boron transporter amplification and GAPDH (NM_008084.2, Forward: 5’-GTG TGA ACG GAT TTG GCC GT-3’, Reverse: 5’-TTG ATG TTA GTG GGG TCT CG-3’) were used as a housekeeping gene. Integrin α_5_ (NM_010577.3, Forward: 5’-GGA CGG AGT CAG TGT GCT G-3’, Reverse: 5’-GAA TCC GGG AGC CTT TGC TG-3’), Integrin β_1_ (NM_010578, Forward: 5’-CAT CCC AAT TGT AGC AGG CG-3’, Reverse: 5’-CGT GTC CCA CTT GGC ATT CAT-3’), Integrin α_v_ (NM_008402.2, Forward: 5’-CAC CAG CAG TCA GAG ATG GA-3’, Reverse: 5’-GAA CAA TAG GCC CAA CGT TC-3’), Integrin β_3_ (NM_016780.2, Forward: GGA ACG GGA CTT TTG AGT GT-3’, Reverse: 5’-ATG GCA GAC ACA CTG GCC AC-3’) and β-actin (NM_007393.3, Forward: 5’-TTC TAC AAT GAG CTG CGT GTG-3’, Reverse: 5’-GGG GTG TTG AAG GTC TCA AA-3’) were used as a housekeeping gene.

The fractional cycle number at which fluorescence passed the threshold (Ct values) was used for quantification by the comparative Ct method. Sample values were normalized to the threshold value of housekeeping gene GAPDH or β-actin: Δ*C*_*T*_ = *C*_*T*_(*experiments*) − *C*_*T*_ (β − actin). The Ct value of the control (B0% substrate) was used as a reference. ΔΔ*C*_*T*_ = Δ*C*_*T*_(*experiment*) − Δ*C*_*T*_(*control*). mRNA expression was calculated by the following equation: 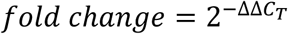

### 5.6. Co-localization experiments

Co-localization of NaBC1/BMPR, NaBC1/α_5_ and α_v_ experiments were performed using DUOLINK® PLA system (Sigma-Aldrich) and following the manufacturer’s instructions. Specific primary antibodies used were: anti-NaBC1 (abcam, 1:200), anti-BMPR1A (abcam, 5 µg mL^-1^), anti-integrin α_5_ (abcam, 1:500) and anti-integrin α_v_ (abcam, 1:500). For image quantification of co-localization fluorescent dots, at least 30 individual cells were imaged for each condition under an epifluorescence microscope (Nikon Eclipse 80i).

### 5.7. In-Cell Western

For evaluation of MLC, ERK, Runx2, FAK, Akt and Smad1 phosphorylation as well as active YAP, NaBC1, integrin α_5_ and α_v_ we used In-Cell Western quantification. MSCs (10.000 cells cm^-2^) were seeded onto FN-coated substrates during 1.5 h at 37 °C and 5% CO_2_. Cells were then fixed using fixative buffer (10 ml formaldehyde, 90 ml PBS, 2 g sucrose) at 37 °C for 15 minutes and then permeabilized in cold methanol at 40 °C for 5 minutes. Cells were then blocked in 0.5% blocking buffer (non-fat dry milk powder in 0.1% PBST buffer) at RT for 2 hours followed by 3 washes of 10 minutes with 0.1% PBST. Cells were then incubated with primary antibodies: pMLC (Cell Signaling, 1:200), Runx2 and pRunx2 (Stratech, 1:100), ERK and pERK (Cell Signaling, 1:200), Smad1 and pSmad1 (Cell Signaling, 1:200), FAK and pFAK (Millipore, 1:200), Akt and pAkt (ThermoFisher, 1:500), active YAP (abcam, 1:500), NaBC1 (abcam, 1:200), BMPR1A (abcam, 5 µg mL^-1^), integrin α_5_ (abcam, 1:500) and anti-integrin α_v_ (abcam, 1:500) diluted in blocking buffer at 4 °C overnight. After 3 washes of 10 minutes with 0.1% PBST buffer, cells were incubated with 1:800 diluted infrared-labeled secondary antibody IRDye 800CW (LI-COR) and 1:500 diluted CellTag 700 Stain (LI-COR) at RT for 1 hour, followed by 5 washes of 10 minutes with 0.1% PBST. Samples were then dried overnight at room temperature. Infrared signal was detected using an Odyssey infrared imaging system.

### 5.8. Image analysis

To analyze focal adhesions, vinculin images were segmented by ImageJ, using Trainable Weka Segmentation plugin to create a binary mask. After segmentation, focal adhesion size and number were determined using different commands of the same software. Values of focal adhesion size frequency were represented using GraphPad Prism 6.0. using a bin width of 0.2 μm. Cell morphology was analyzed by calculation of different parameters using ImageJ software. Staining intensity of immunofluorescence images were quantified by ImageJ software.

### 5.9. Statistical Analysis

For statistical analysis, the data was analyzed for normality using the D’Agostino and Pearson omnibus normality test with an alpha of 0.05. When the normality test was passed, an ordinary one-way ANOVA test with a Tukey’s, Sidak’s or Dunnett’s post-hoc analysis (p = 0.05) was used to compare the means of the columns against the control column. When the normality test was not passed, a non-parametric test with a post-hoc Dunn analysis (p = 0.05) was used to compare the means of each column against the control column. Data is represented as mean ± standard deviation. Sample size for each statistical analysis is indicated in the corresponding figure legends. GraphPad Prism 6 XML software has been used for statistical analysis.

## Supporting Information

Supporting Information is available from the Wiley Online Library or from the author.

## Supplementary information

**Figure S1.**
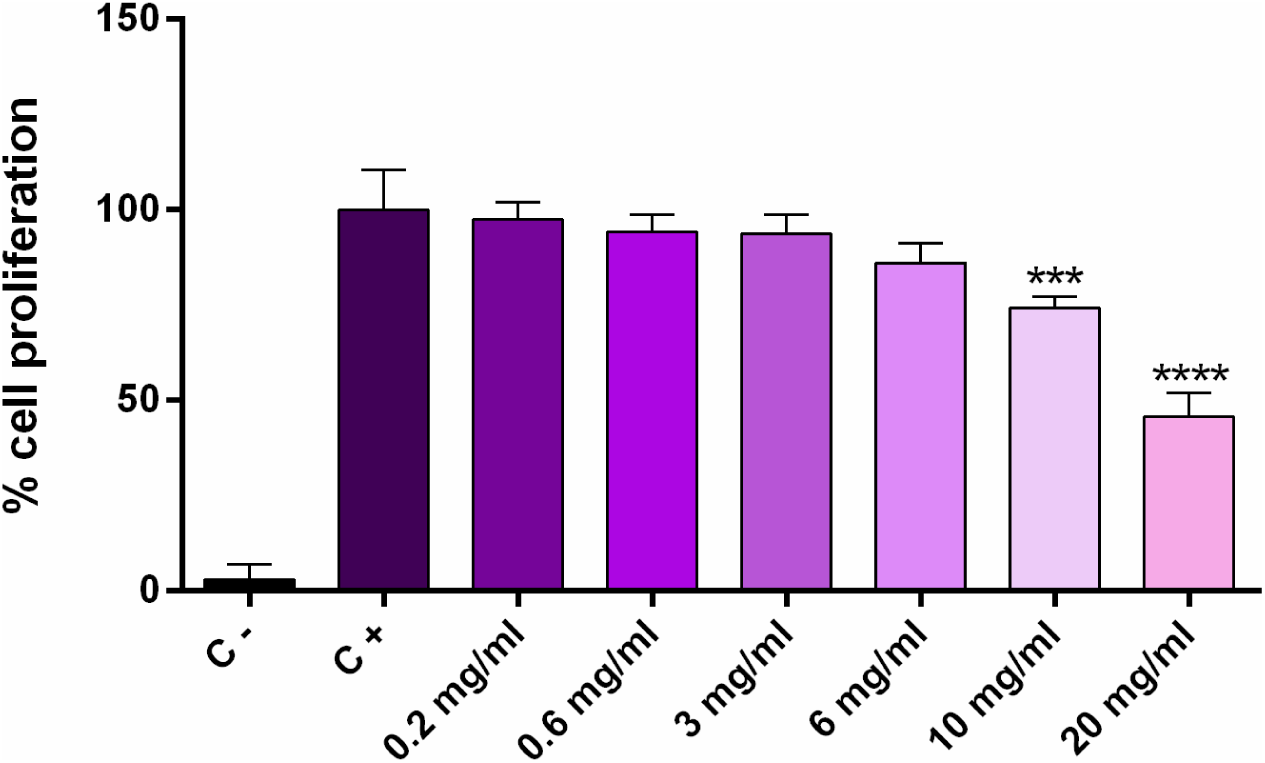
Cytotoxicity of boron. Results of MTS assay after 24 h of culture. 20.000 cells cm^-2^ of MSCs were seeded and their metabolic activity was measured after 24 h of incubation with different quantities of boron in the medium (0.2, 0.6, 3, 6, 10 and 20 mg mL^-1^). The formation of formazan product was followed by measuring absorbance at 490 nm. All measurements were performed in triplicate. Statistics are shown as mean ± standard deviation. Data was analyzed by an ordinary one-way ANOVA test and corrected for multiple comparisons using Tukey’s analysis (P = 0.05). ***p < 0.001, ****p < 0.0001.

**Figure S2.**
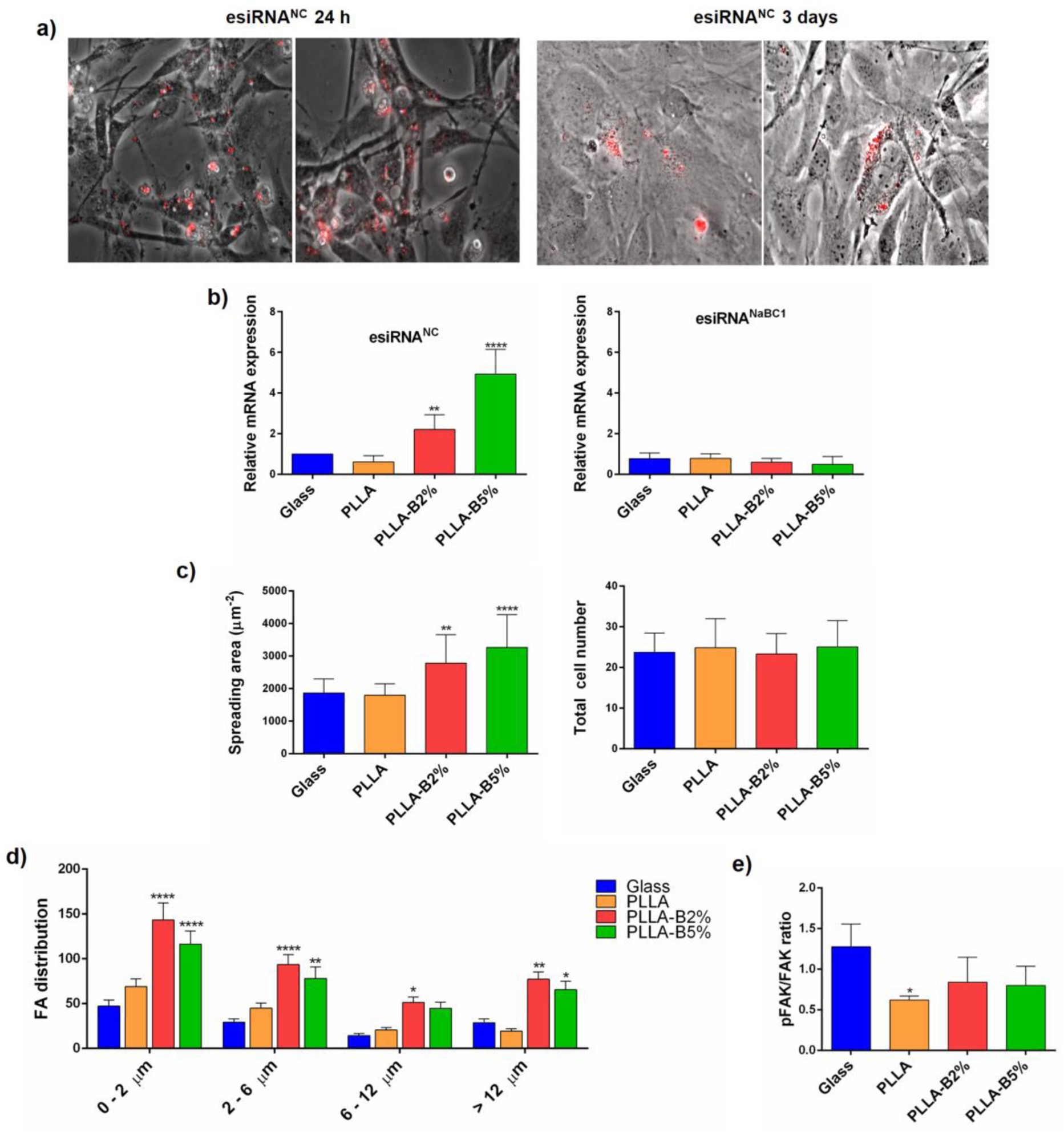
Boron effects on focal adhesion (FA). a) Microscopic images obtained after 24 h and 3 days of culture of transfected MSC cells with a universal negative control (esiRNA^NC^) labelled with Cyanine 3 as a control for assessment of silencing efficiency. b) qPCR amplification of NaBC1 mRNAs from transfected MSCs with esiRNA^NC^ and esiRNA^NaBC1^. esiRNA^NaBC1^ transfected cells express reduced NaBC1 mRNAs confirming an efficient silencing of boron transporter. n = 6 different biological replicas. c) Image analysis quantification of different parameters related to cell adhesion. Spreading area and total cell number. Boron induces higher cell spreading. n = 20 images/condition from three different biological replicas. d) Focal adhesion distribution. Boron induce higher levels of nascent (0-6 µm) and mature FA (6-12 µm) formation. e) In-Cell Western quantification of pFAK/FAK ratio. Boron-induced FA formation is not dependent on FAK phosphorylation. n = 4 different biological replicas. Statistics are shown as mean ± standard deviation. Data was analyzed by an ordinary one-way ANOVA test and corrected for multiple comparisons using Dunnett analysis (P = 0.05). *p < 0.05, **p < 0.01, ***p < 0.001, ****p < 0.0001.

**Figure S3:**
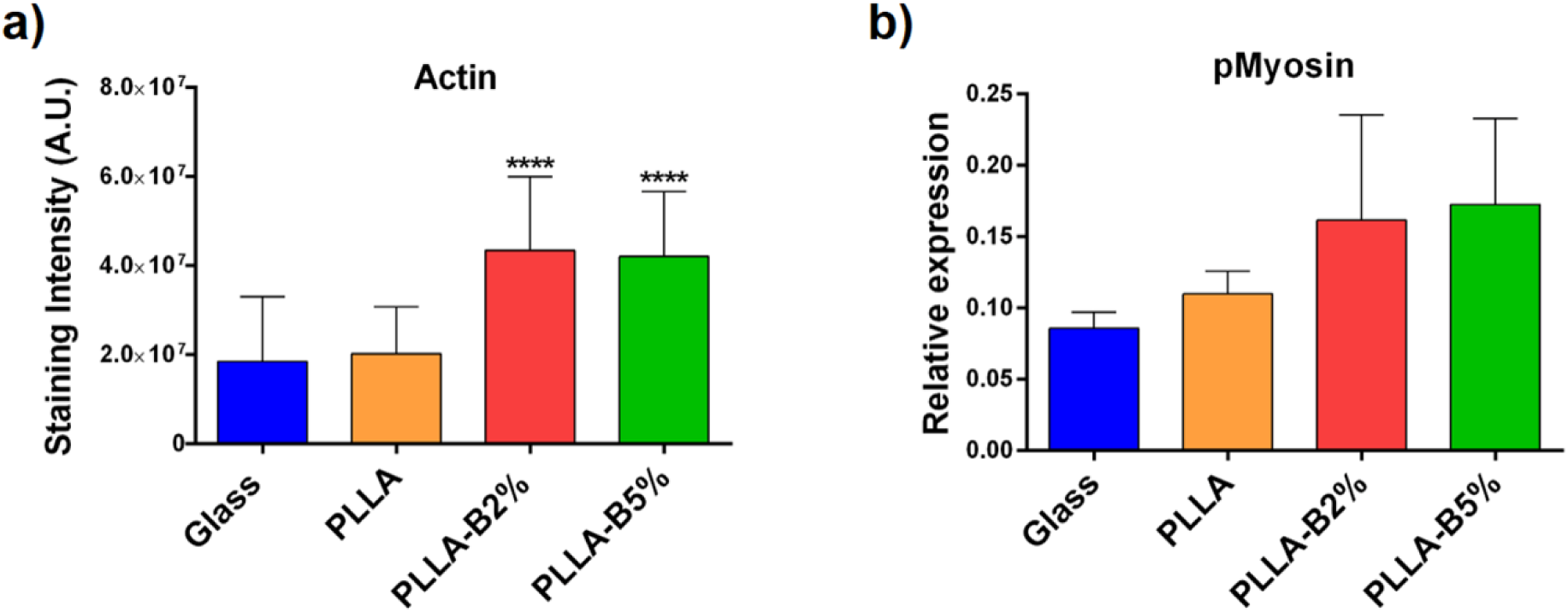
Boron effects on myosin light chain phosphorylation and actin stress fiber formation. a) Image analysis quantification from images of Figure 2, of actin fibers as a parameter related to cell contractility. Active NaBC1 induces cell contractility. n = 20 images/condition from three different biological replicas. b) In-Cell Western quantification of pMyosin. Active NaBC1 induce myosin light chain phosphorylation. n = 4 different biological replicas. Statistics are shown as mean ± standard deviation. Data was analyzed by an ordinary one-way ANOVA test and corrected for multiple comparisons using Dunnett analysis (P = 0.05). ****p < 0.0001.

**Figure S4:**
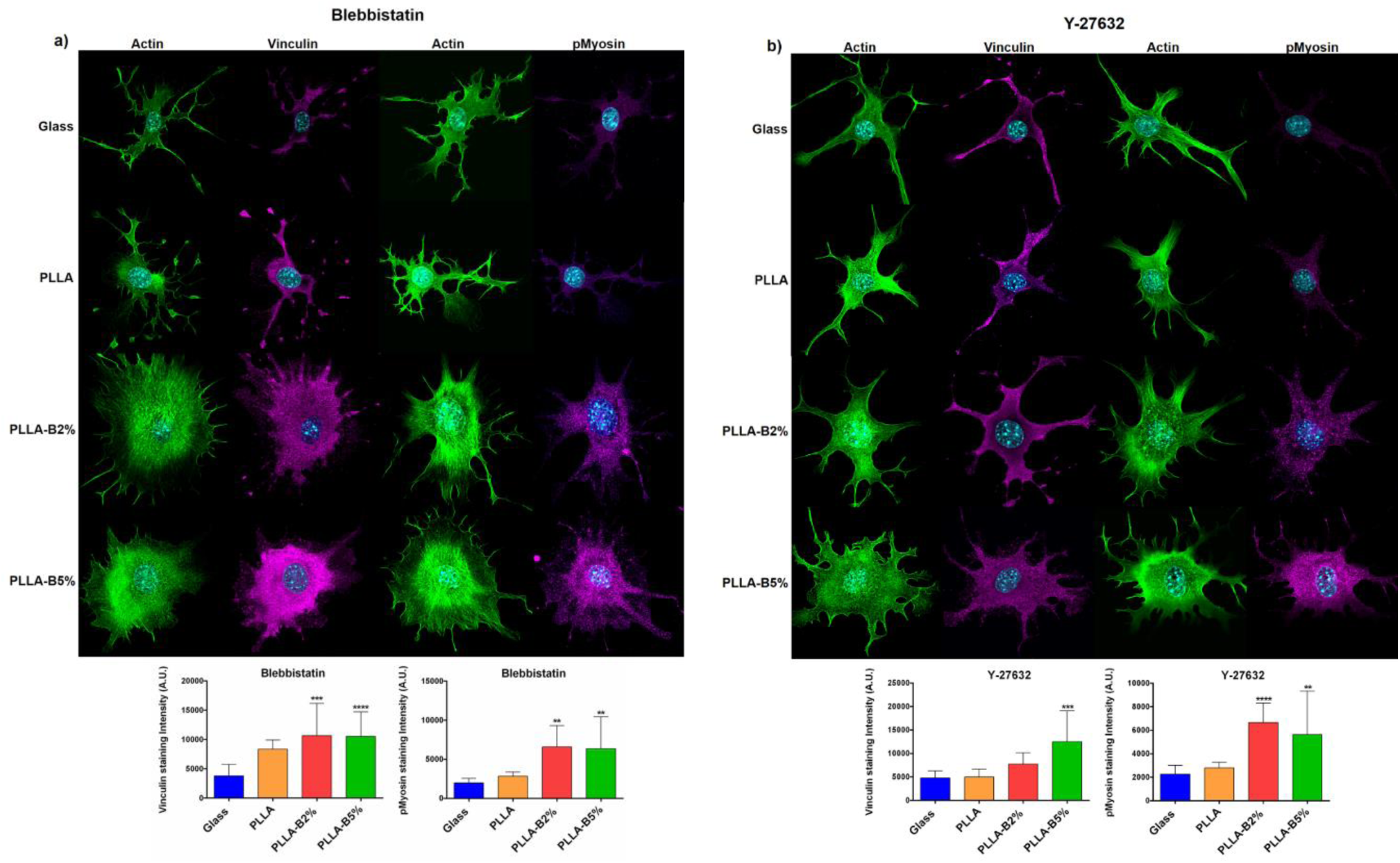
Analysis of contractility using pharmacological inhibitors. a) Immunofluorescence images of actin cytoskeleton (green), nuclei (cyan), vinculin and pMLC (pMyosin, magenta) after inhibition of contractility with Blebbistatin (Myosin II inhibitor). The PLLA-B2% and PLLA-B5% substrates presented higher levels of vinculin and pMyosin staining even after Blebbistatin addition compared to PLLA and Glass control substrates. n = 20 images/condition from three different biological replicas. b) Immunofluorescence images of actin cytoskeleton (green), nuclei (cyan), vinculin and pMLC (pMyosin, magenta) after inhibition of contractility with Y-27632 (Rho-kinase inhibitor). The PLLA-B2% and PLLA-B5% substrates presented higher levels of vinculin and pMyosin staining even after Y-27632 addition compared to PLLA and Glass control substrates. n = 20 images/condition from three different biological replicas. Statistics are shown as mean ± standard deviation. Data was analyzed by an ordinary one-way ANOVA test and corrected for multiple comparisons using Dunnett analysis (P = 0.05). **p < 0.1, ***p <0.001, ****p < 0.0001.

**Figure S5.**
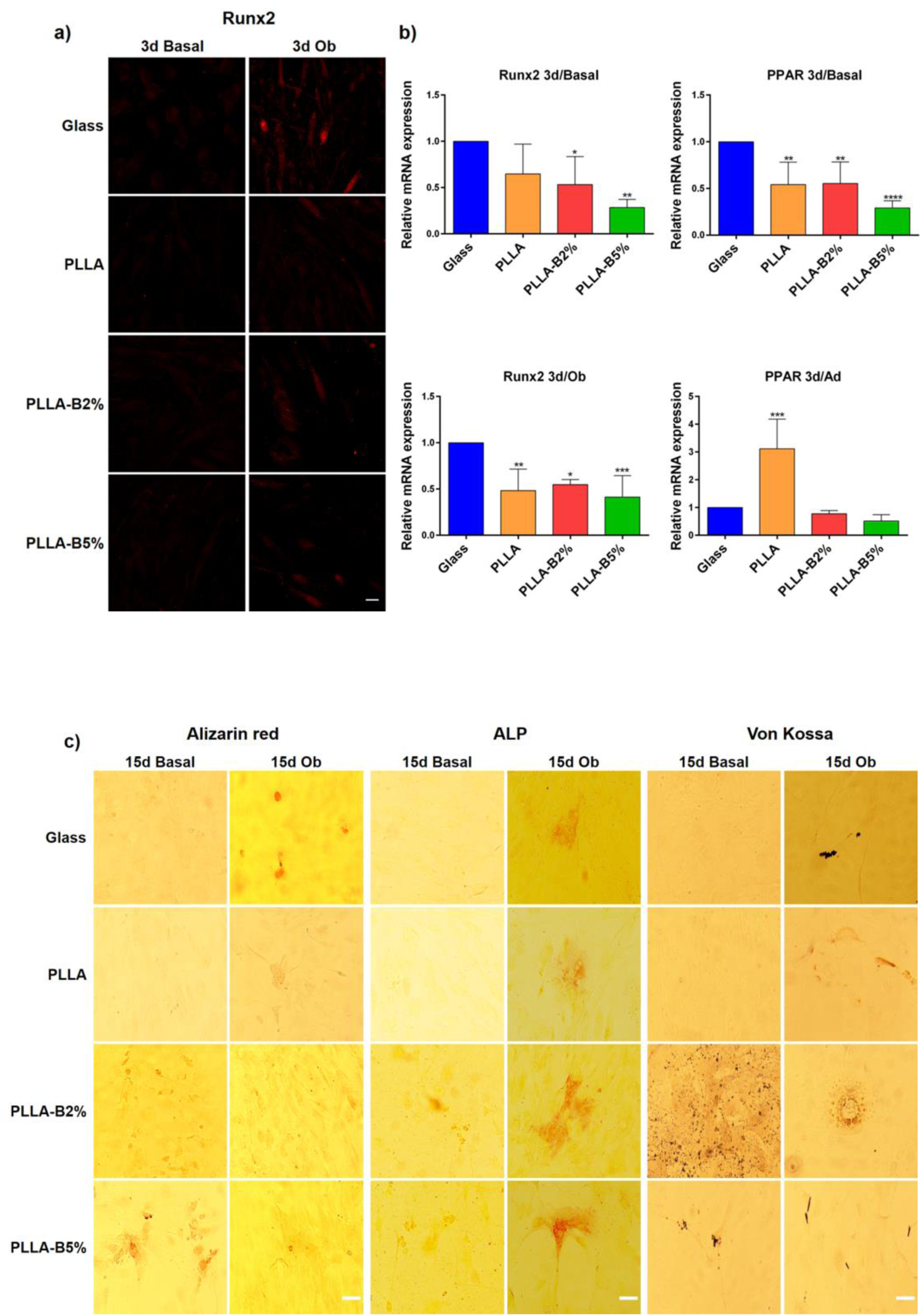
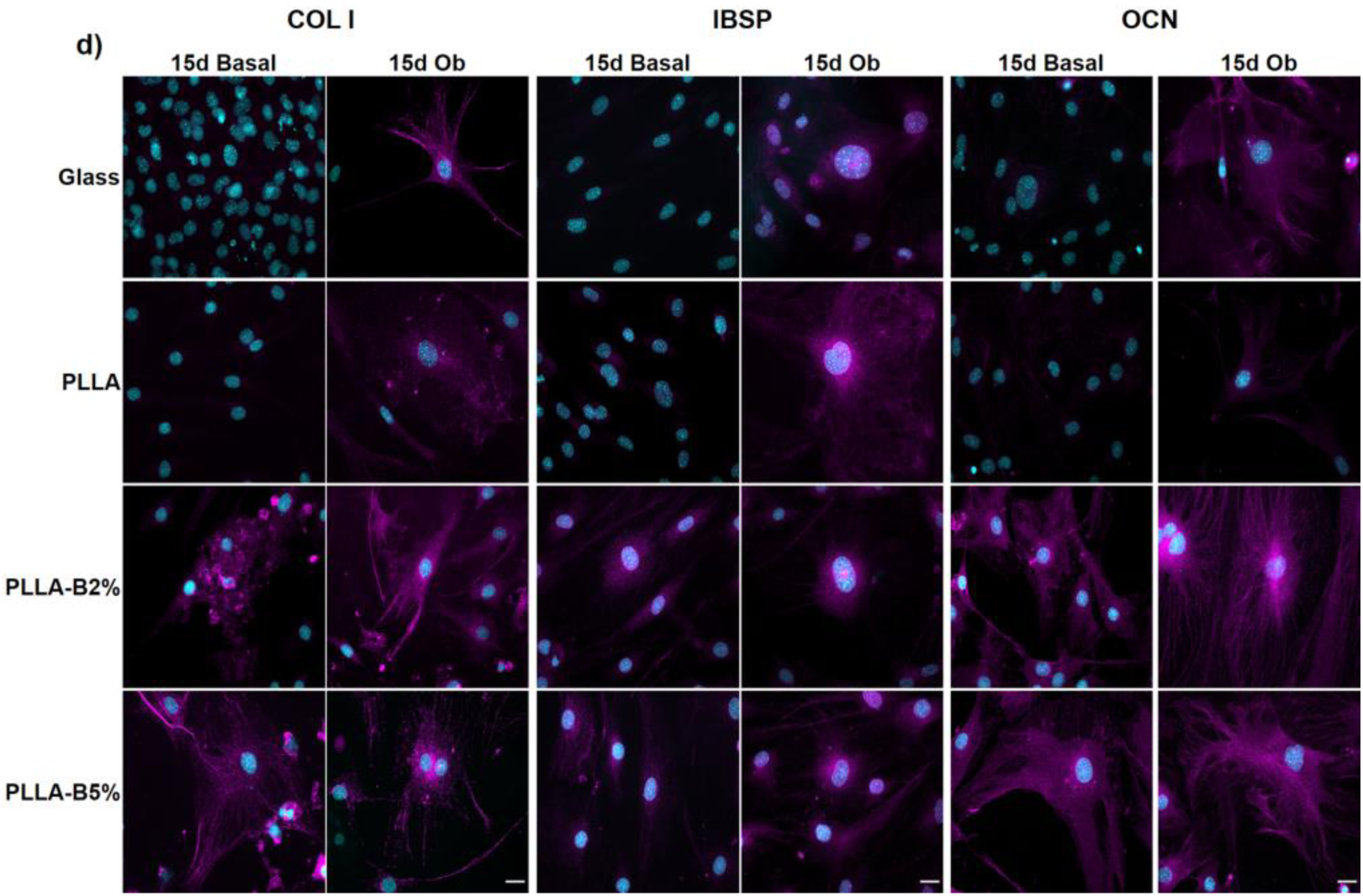
Boron effects on MSC differentiation. a) Immunofluorescence images of MSCs cultured for 3 days onto functionalized substrates (FN-coated) and boron (PLLA-B2%, PLLA-B5%) in culture medium, under basal and differentiation conditions (osteogenic), for detection of early osteogenic marker (Runx2, red). Scale bar 25 µm. b) qPCR analysis of relative mRNA expression of early expressed transcription factors involved in osteogenic (Runx2) and adipogenic (PPARγ2) lineage determination. RNA was extracted after 3 days of culture under both basal and differentiation conditions (osteogenic and adipogenic differentiation media). n = 4 different biological replicas. c) Histological staining of MSCs cultured for 15 days onto functionalized substrates (FN-coated) and boron (PLLA-B2%, PLLA-B5%) in culture medium, under basal and differentiation conditions (osteogenic), for detection of late osteogenic markers (Alizarin Red-calcium deposits in red, Alkaline Phosphatase activity (ALP) in red and Von Kossa-calcium deposits in black). Scale bar 25 µm. d) Immunofluorescence images of MSCs cultured for 15 days onto functionalized substrates (FN-coated) and boron (PLLA-B2%, PLLA-B5%) in culture medium, under basal and differentiation conditions (osteogenic), for detection of late osteogenic markers: Collagen I (Col I), Integrin binding sialoprotein (IBSP) and Osetocalcin (OCN). Scale bar 25 µm. B alone induces osteogenic markers expression. Statistics are shown as mean ± standard deviation. Data was analyzed by an ordinary one-way ANOVA test and corrected for multiple comparisons using Dunnett analysis (P = 0.05). *p < 0.05, **p < 0.01, ***p <0.001, ****p < 0.0001.

**Figure S6.**
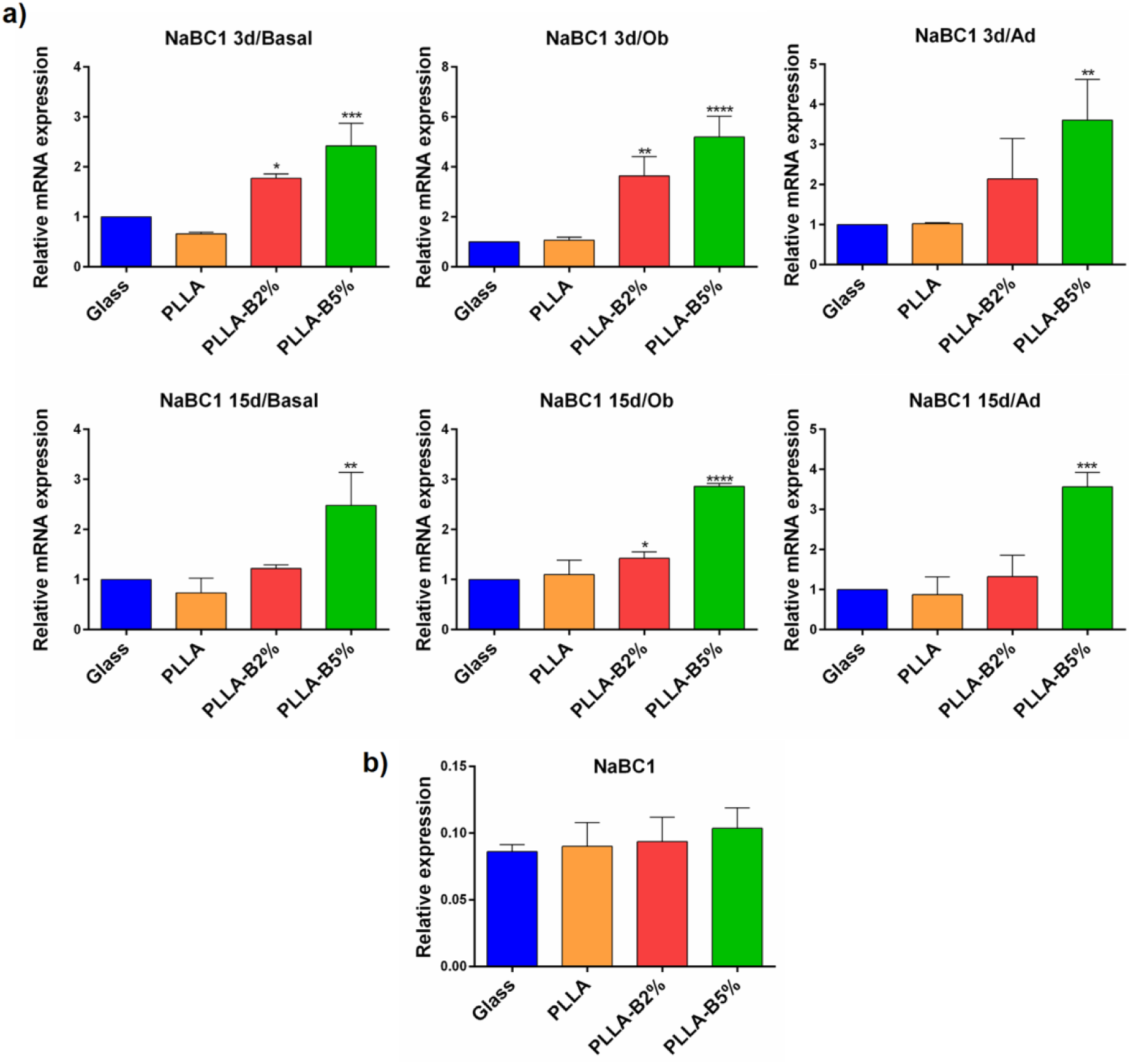
NaBC1 gene expression. a) qPCR analysis of relative mRNA expression of boron transporter NaBC1. RNA was extracted from MSCs after 3 or 15 days of culture under both basal and differentiation conditions (osteogenic and adipogenic differentiation media). n = 5 different biological replicas. b) In-Cell Western quantification of NaBC1. n = 4 different biological replicas. Statistics are shown as mean ± standard deviation. Data was analyzed by an ordinary one-way ANOVA test and corrected for multiple comparisons using Tukey analysis (P = 0.05). *p < 0.05, **p < 0.01, ***p < 0.001, ****p <0.0001.

**Figure S7.**
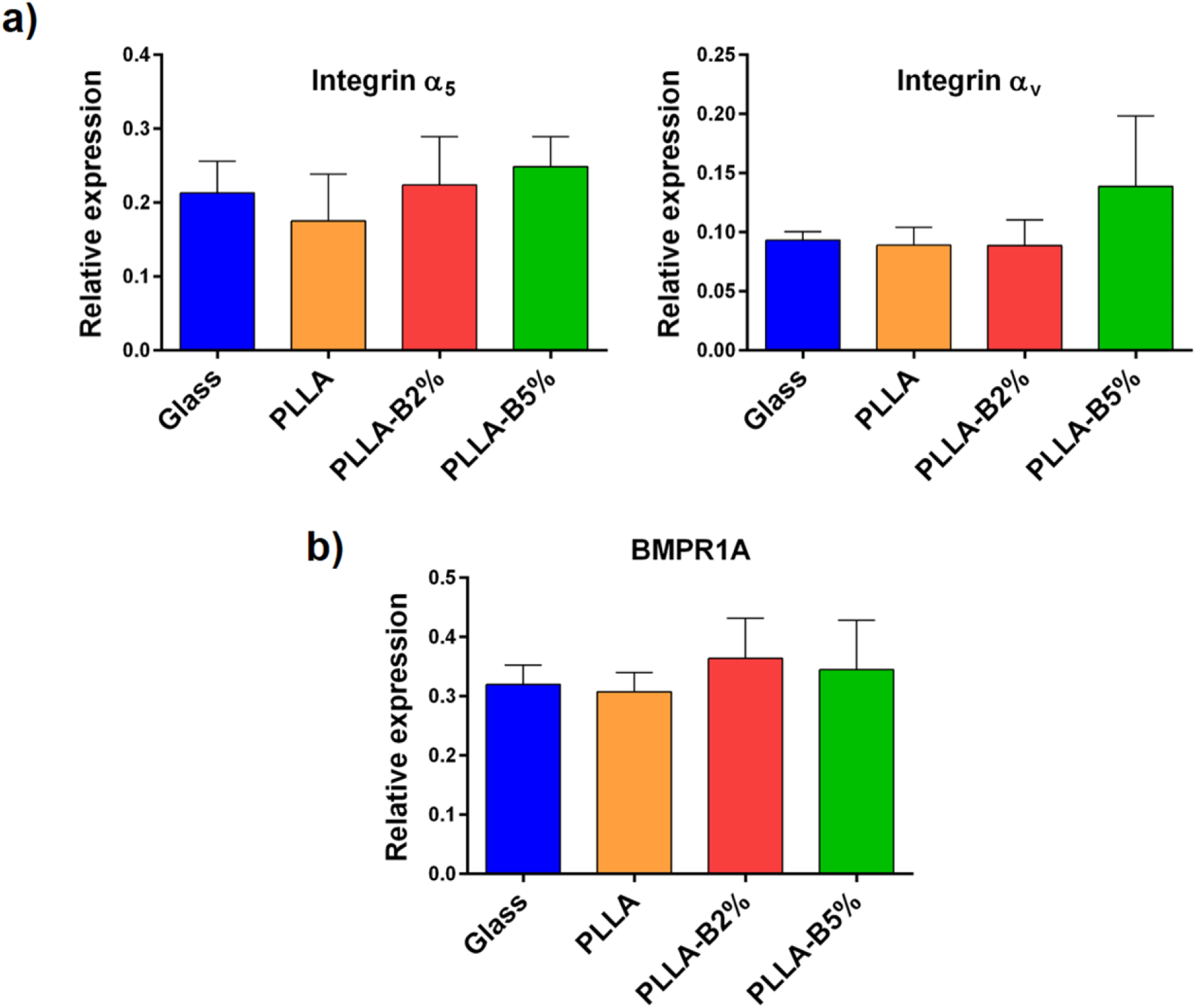
In-Cell Western quantification of FN-binding integrins and BMPR1A. a) In-Cell Western quantification of α_5,_ α_v_ integrins. n = 4 different biological replicas. b) In-Cell Western quantification of BMPR1A. n = 4 different biological replicas.

